# Gene flux shapes diversity and evolution of the ancient 17q21.31 inversion polymorphism

**DOI:** 10.64898/2026.09.28.755014

**Authors:** Olivia S. Harringmeyer, Arjun Biddanda, Rajiv C. McCoy, Joshua M. Akey

## Abstract

A hallmark of chromosomal inversions is that they suppress recombination between haplotypes, allowing inversion haplotypes to persist as single co-inherited units. To determine the extent to which inversions nevertheless permit genetic exchange, we investigated a common 979-kb inversion polymorphism at the human 17q21.31 locus. This locus exhibits deep divergence between the reference (H1) and inverted (H2) haplotypes, extensive segmental duplications (SDs) flanking the inversion, and association with neurodegenerative diseases, developmental disorders, and fertility-related phenotypes. Using single-cell sperm genome sequencing data, we directly measured recombination rates between H1 and H2 haplotypes and found near-complete suppression of single-crossover events between the haplotypes. The rare single crossovers that did occur were mediated by non-allelic homologous recombination between shared H1 and H2 SDs, generating novel duplication architectures. In contrast, two-switch events consistent with gene conversion or double crossovers, spanning 17-150 kb, occurred throughout the inversion at rates exceeding genome-wide estimates for events of comparable size. Consistent with recurring genetic exchange, we identified 99 distinct H1-H2 recombinant haplotypes segregating in All of Us genomes, including 26 with combinations of H1 and H2 SDs. These recombinant haplotypes facilitated dissection of the inversion’s effects on fertility-related phenotypes; using a large parent-embryo dataset, we found that H2 additively increases female crossover rates across chromosomes and that *KANSL1* duplications do not explain this effect. Finally, ancestral recombination graphs dated H1-H2 gene flux (the exchange of genetic material between alternative arrangements) to approximately 100-500 thousand years ago, revealing that H1 and H2 haplotypes have co-segregated for at least half a million years. Together, these results demonstrate that inversions can be permeable barriers to recombination, with ongoing gene flux influencing the diversity and evolution of inversion polymorphisms.

## Introduction

Chromosomal inversions were one of the first examples of genetic polymorphisms, initially discovered in *Drosophila*^1,2^, and have since been found within diverse plant and animal species^3^. Inversions are a unique mutation type that does not change DNA content but alters the orientation of a chromosomal segment. Inversions affect the genome in two ways: they disrupt sequence at inversion breakpoints and suppress recombination within inverted sequences^4^. These properties allow inversions to reshape patterns of genetic variation and inheritance, yet how breakpoint architecture and recombination suppression interact to influence the evolution of inversion polymorphisms remains poorly understood.

Inversion breakpoints are often repetitive and highly complex genomic regions. The breakpoints of large inversions in mammals are enriched for segmental duplications (SDs) – long (>1-kb) low-copy number repeats that drive the formation of inversions via ectopic recombination^5–7^. Such duplication architectures of inversion breakpoints make them prone to recurrent inversion and other complex structural variation^8,9^. Inversion breakpoints can influence phenotypes through disrupting gene sequence and expression or driving additional mutation^7,10^. Whether complex breakpoint architectures also influence genetic exchange between alternative inversion arrangements remains largely unexplored.

Inversions are also modifiers of recombination. They suppress effective recombination between arrangements because single crossovers within inversion heterozygotes can generate gametes with unbalanced recombinant chromosomes^3^. By suppressing recombination, inversions keep mutations in tight linkage disequilibrium (LD), including mutations captured by or subsequently accumulated within the inversion^11^ and potentially beyond its breakpoints^12^. Inversions often form the structural basis for supergenes, in which sets of alleles are co-inherited as single genetic units^13^. Likely through this effect, inversions contribute to adaptation^14–17^, sex chromosome evolution^18–20^ and speciation^21,22^.

Recombination suppression is thus a defining feature of inversions, but the extent to which inversions limit gene flux between alternative arrangements remains poorly understood. How much genetic exchange occurs between inverted and standard haplotypes, and through which mechanisms^23^, has rarely been measured (but see ^24,25^), especially in humans. In the absence of gene flux, alternative arrangements may evolve as largely independent lineages. In contrast, recurrent genetic exchange can generate haplotype diversity, break down linkage among variants within inversions, and potentially reduce the deleterious consequences of suppressed recombination^26^, thereby influencing inversion function and evolution. Resolving the extent, mechanisms, and evolutionary consequences of gene flux is therefore important for understanding how inversion polymorphisms evolve and persist^3,27,28^.

The 17q21.31 inversion polymorphism, one of the most common inversions in the human genome, provides a rare opportunity to dissect both breakpoint and recombination effects of an inversion.

The 979-kb inversion (referred to as H2) is segregating in many human populations, with frequencies reaching ~20% in Europe, and is strongly differentiated from the reference haplotype (H1)^29^, with the age of the inversion estimated at >2 Mya^30^. H2 shows signatures of recent, positive selection in Europe; the association of H2 with increased maternal crossover rates and offspring count likely contributes to its positive selection^29^. H2 is also associated with protection from neurodegenerative diseases (Parkinson’s, progressive supranuclear palsy, corticobasal degeneration)^31^. The inversion is flanked by large, complex duplications that make 17q21.31 susceptible to additional structural mutation^30,32–34^, including a microdeletion syndrome (Koolen-de Vries)^35^. Using sperm genome sequencing, biobank-scale analyses of recombination phenotypes, and ancestral recombination graphs, we measure recombination between H1 and H2 haplotypes and uncover interconnected effects of the 17q21.31 inversion’s breakpoints and modification of recombination, which combine to shape the evolutionary history of this longstanding polymorphism.

## Results

### Structural variation at the 17q21.31 locus

The 17q21.31 locus is a complex genomic region with a 979-kb inversion. The inverted region has a 478-kb stretch of single-copy sequence, and the rest consists of segmental duplications (SDs), with H1 denoting the reference genome orientation (CHM13 and GRCh38) and H2 the inverted orientation (Figure 1A). Each orientation consists of several subhaplogroups defined by the number of copies of α (146-kb partial duplication of *KANSL1*), β (241-kb partial duplication of *KANSL1* and *LRRC37A*) and γ (197-kb duplication of *LRRC37A* and *NSF*) SDs: H1 haplotypes contain 1-3 copies of β and 1-4 copies of γ whereas H2 haplotypes contain 1-2 copies of α and 1-2 copies of γ^30,32,33^. Representative haplotypes from the Human Pangenome Reference Consortium^36^ and Human Genome Structural Variation Consortium^37^ show the location and orientation of these SDs in different subhaplogroups; β duplications are tandem in H1 haplotypes whereas α duplications flank the inverted region in H2 haplotypes (Figure 1B). Both CHM13 and GRCh38 represent subhaplogroup H1.β1.γ2 (Figure 1B,C).

**Figure 1.**
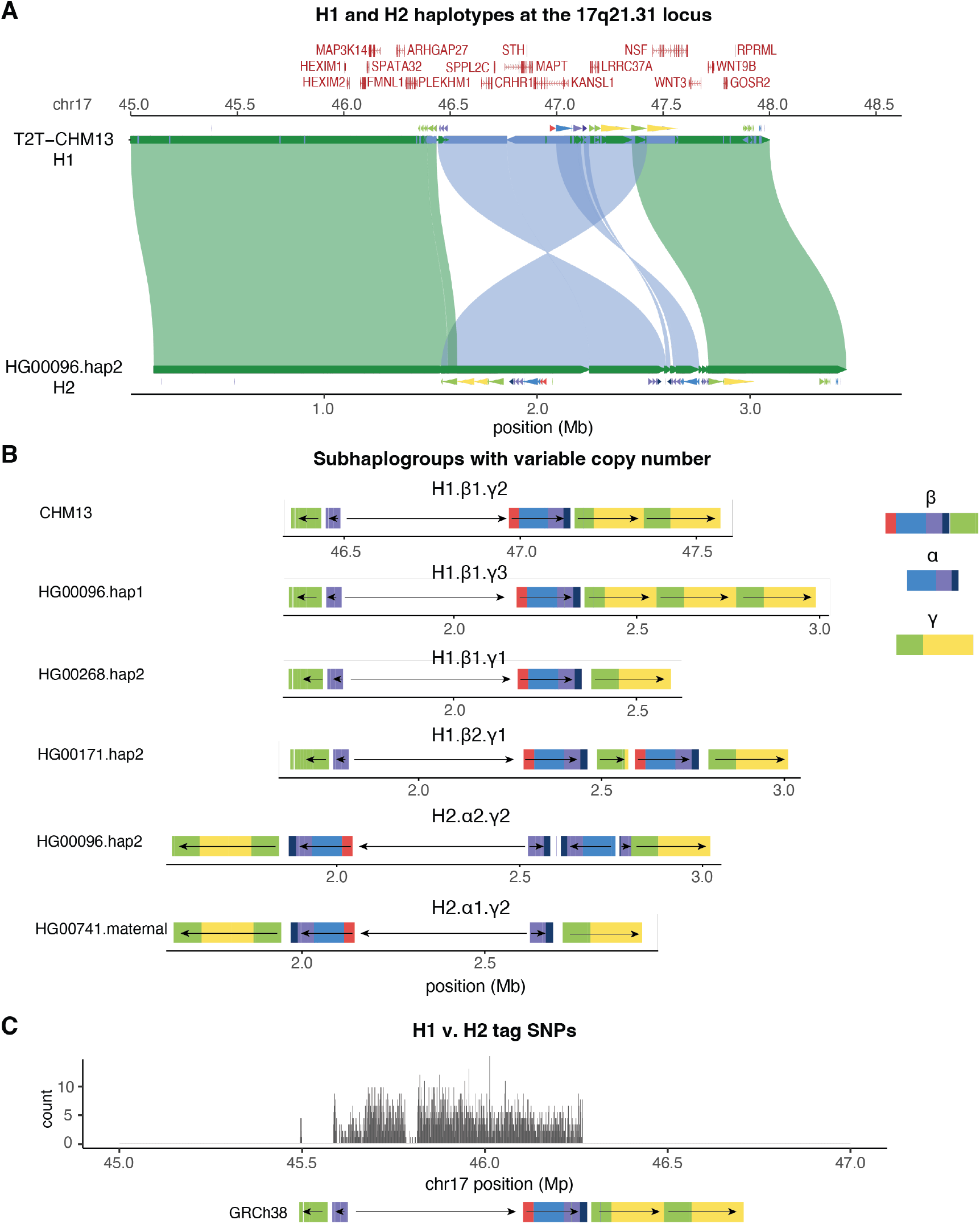
An overview of the 17q21.31 locus and its structural diversity. **A)** Alignments (>200-kb) between T2T-CHM13 reference genome, which harbors the H1 haplotype, and a long-read haplotype-resolved genome assembly harboring the H2 haplotype, visualized with SVbyEye. SDs are shown with arrows, and genes are shown above. **B)** Representative long-read haplotypes from HPRC and HGSVC3 for subhaplogroups at the 17q21.31 locus. Subhaplogroups are defined by copy number at the α, β and γ regions, with SDs colored by family. **C)** The location and count of the 2,461 H1 v. H2 tag SNPs along GRCh38, with SD locations in GRCh38 shown below.

High divergence between H1 and H2 haplotypes facilitates genotyping the inversion with short-read sequencing data. Genetic variation within the inversion region separates samples into three distinct PCA clusters corresponding to the three inversion genotypes (H1/H1, H1/H2 and H2/H2) (Supplemental Figure 1). We identified 2,461 SNPs that tag H1 versus H2 haplotypes (>95% frequency in H2/H2, <5% frequency in H1/H1) in the 1000 Genomes Project (1KGP) dataset; these SNPs span the single-copy region of the inversion as well as the *KANSL1* SDs (α and β) (Figure 1C), facilitating subhaplogroup analyses in short-read sequencing datasets.

### H1-H2 recombination in the male germline

While it is well-established that inversions suppress recombination in heterozygotes, direct estimates of recombination rates between inversion and standard haplotypes remain rare. To quantify recombination rates between H1 and H2, we used single-cell whole-genome sperm sequencing data from 20 donors^38^. Genotyping these donors using the H1-H2 tag SNPs, we identified 5 donors with H1/H2 and 15 with H1/H1 inversion genotypes. We inferred crossover events in 974 - 2,274 sperm cells/donor between maternal and paternal haplotypes of each donor (see Methods). H1/H2 donors show significantly lower recombination rates (for single crossovers) within the inversion region than H1/H1 donors (0.09 cM/Mb vs. 0.26 cM/Mb, respectively; permutation test, *p* < 0.01) (Figure 2A). Indeed, in H1/H2 donors, there were no single crossover events found within the single-copy region of the inversion (Figure 2B). However, when including the full inversion region (flanking SDs +/− 500-kb), the single crossover rate between H1 and H2 haplotypes was non-zero. We identified 17 sperm cells with single crossover events between H1 and H2 and localized most of these events to the SDs at the inversion breakpoints (Figure 2B, Supplemental Table 1). We note that these crossover events occurred in both directions, from H1 to H2 and from H2 to H1 (Figure 2B). Based on subhaplogroup assignments for the H1/H2 sperm donors (Supplemental Figure 2), we found that the single-crossover events in H1/H2 sperm likely occurred as non-allelic homologous recombination (NAHR) between colinear SDs found on H1.β1.γ1 (or H1.β1.γ2 or H1.β2.γ1) and H2.α2.γ2 haplotypes (Figure 2C). These results suggest that the SD architecture of H1 and H2 haplotypes, in which high identity duplications flanking the inversion are colinear between haplotypes, promotes crossover events between H1 and H2.

**Figure 2.**
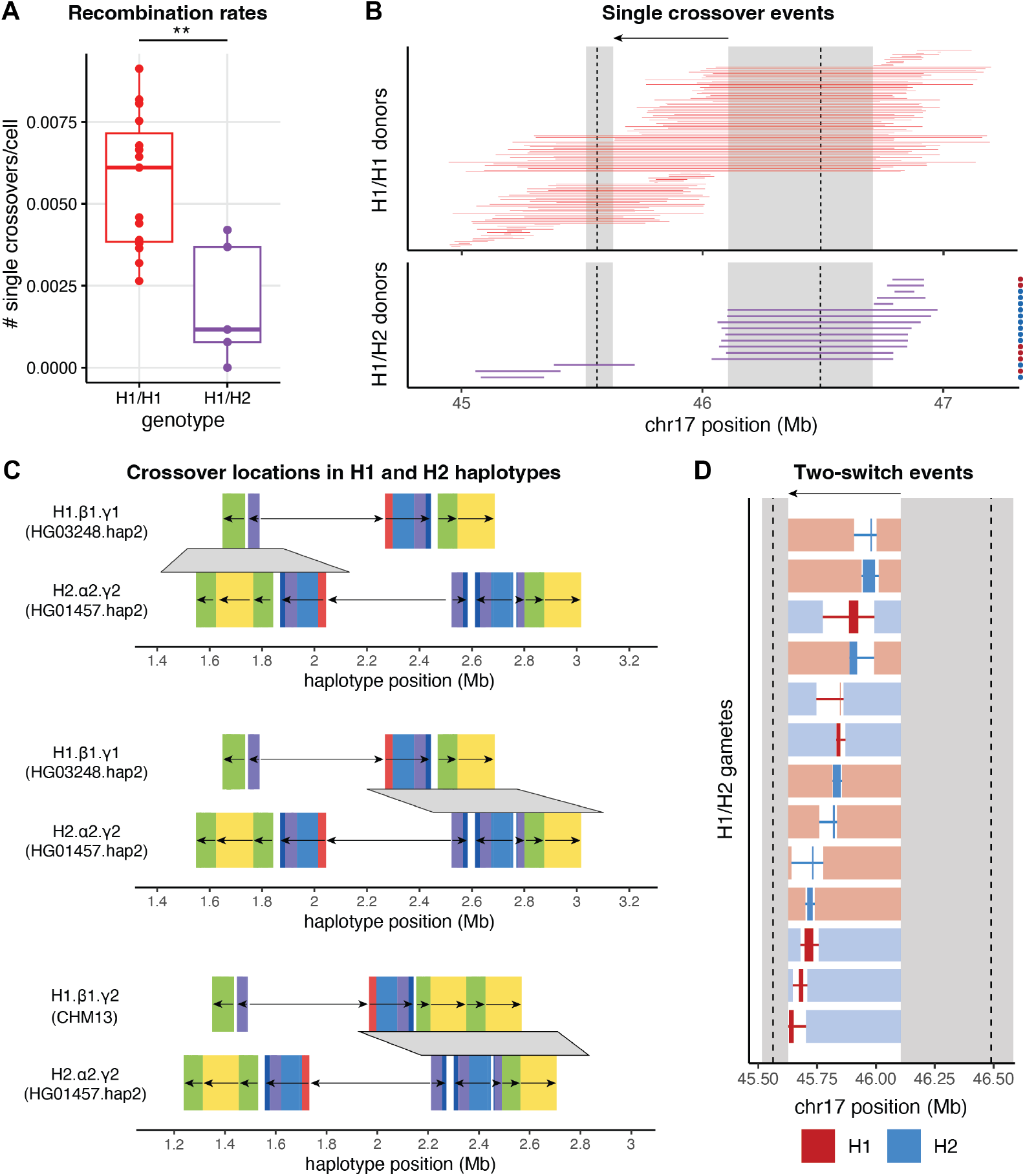
H1-H2 recombination in single-cell sperm sequencing. **A)** Rates of single-crossovers within the inversion region +/− 500-kb, comparing H1/H1 sperm donors (*n*=15) and H1/H2 sperm donors (*n*=5) (*p*<0.01, permutation test (see Methods)). **B)** Locations of single crossover events, shown as interval between the two defining heterozygous SNPs, in sperm from H1/H1 donors (top, red) and H1/H2 donors (bottom, purple). Inversion region (single-copy) is shown with the black arrow, with flanking SD regions highlighted in gray. For sperm cells from H1/H2 donors, background haplotype is denoted with red dot for H1 and blue dot for H2 based on genotypes within the single-copy region of the inversion. **C)** Three example H1-H2 single crossover events projected onto long-read haplotype-resolved genome assemblies that match subhaplogroups for the H1/H2 sperm donors. SDs are shown as colored blocks, with gray regions representing intervals in which crossover events occurred. **D)** Locations of two-switch events in sperm cells from H1/H2 donors. The crossover defining intervals are shown as thin lines, with colors indicating H1 (red) and H2 (blue) haplotypes. The inversion region (single-copy) is shown with the black arrow, and flanking SDs are highlighted in gray.

We also characterized two-switch events, consistent with gene conversion or double crossovers, within the inversion region between H1 and H2 haplotypes, facilitated by the high SNP density between H1 and H2. We identified 13 sperm cells with two-switch tracts within the inversion region, occurring in each of the 5 H1/H2 donors and ranging from 17- to 150-kb in size (Figure 2D, Supplemental Table 1) (note that we could not detect smaller tracts < 5-kb due to the low-coverage sperm data). We found that these events occur across the single-copy region of the inversion and on both H1 and H2 haplotypes (Figure 2D). The rate for such two-switch events (>10-kb) between H1 and H2 haplotypes is 0.0033±0.0008 events per Mb per meiosis, higher than genome-wide rates of gene conversion events >10-kb estimated as 6×10^−5^ or 2.6×10^−4^ events per Mb per meiosis^39,40^. To explore the likely mechanism underlying the observed two-switch events, we compared the per-generation expected number of gene conversion versus double crossover events of this size (>5-kb and <250-kb) across a sequence of length similar to the inversion. Based on recombination rate, crossover interference, and gene conversion tract length estimates^39,41^, we predict that two-switch tract lengths of this size are over 1,000 times more likely to be associated with gene conversion than double crossover events (Supplemental Figure 3).

### Recombinant haplotypes in the All of Us dataset

To explore H1-H2 recombinant haplotypes segregating in humans, we analyzed the All of Us dataset which contains short-read whole-genome sequencing for 414,830 individuals. Using H1-H2 tag SNPs to genotype the 17q21.31 inversion, we found that 69.1% of All of Us individuals were H1/H1, 25.9% H1/H2 and 3.3% H2/H2 (Figure 3A). We assigned copy number at α and β regions using read depth (see Methods): diploid copy number of β ranges from 2-5 in H1/H1 and α ranges from 2-4 in H2/H2 samples (Figure 3B). In H1/H2 samples, we used allelic balance at tag SNPs spanning α and β regions to assign subhaplogroup to each haplotype (Supplemental Figure 4) and found that subhaplogroup H1.β1 is most common (76.7%) followed by H1.β2 (22.4%) and H1.β3 (0.9%) amongst H1 haplotypes and H2.α2 is most common (91.7%) followed by H2.α1 (8.0%) amongst H2 haplotypes (Figure 3B), consistent with previous characterizations of subhaplogroup frequencies^32,33^. We also identified H1/H2 individuals harboring 4 H1 β copies (and 1-2 H2 α copies) and individuals harboring 3 H2 α copies (and 1-2 H1 β copies), indicative of rare H1.β4 (0.03%) and H2.α3 (0.25%) haplotypes respectively (Supplemental Figure 4); however, since the data are unphased, these may also represent H1-H2 recombinants involving α and β regions (see below). These results hinted at the presence of rare, previously undescribed 17q21.31 subhaplogroups segregating in the All of Us genomes.

**Figure 3.**
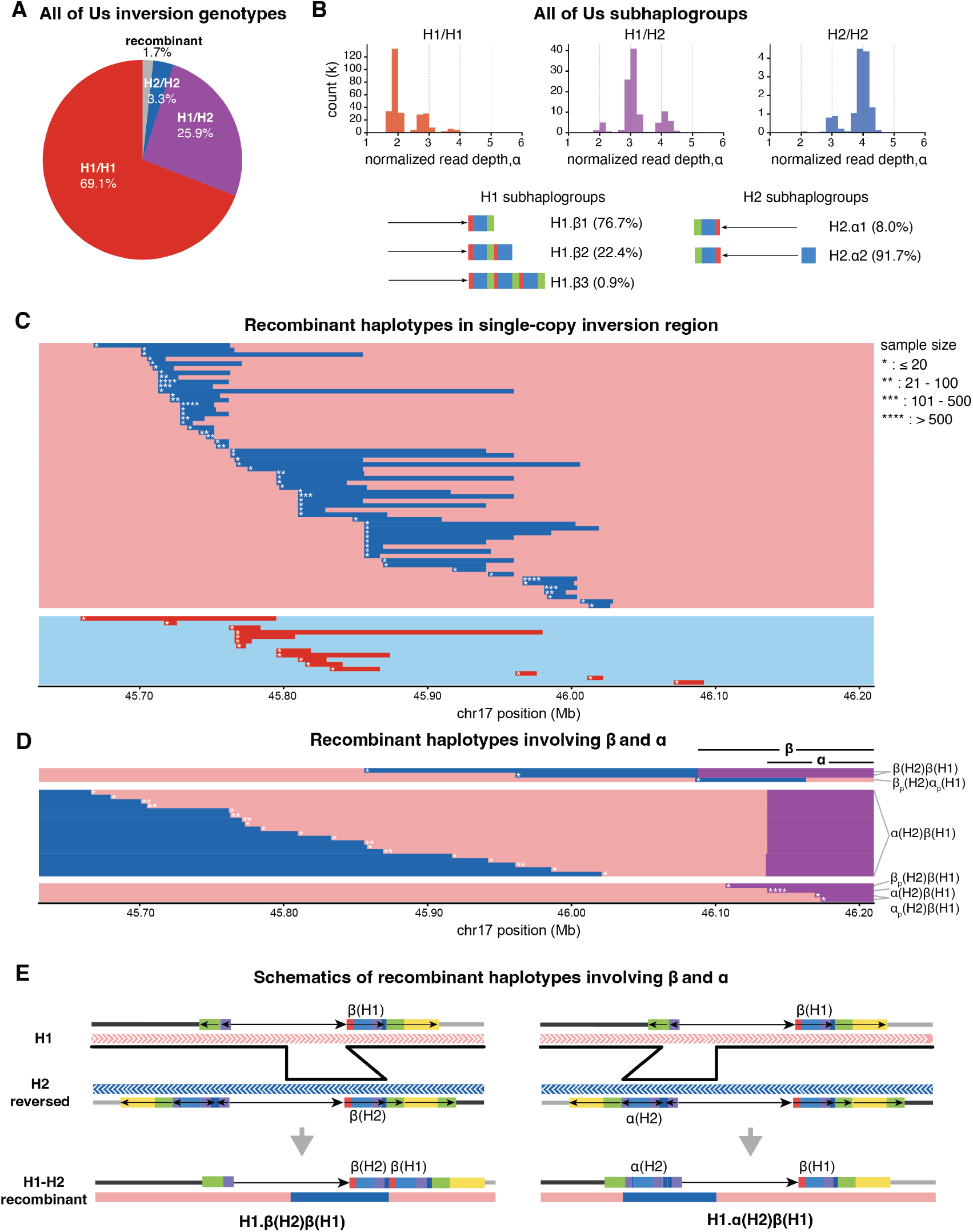
H1-H2 recombinant haplotypes in the All of Us dataset. **A)** Inversion genotypes for the 414,830 All of Us samples with short-read whole-genome sequencing. **B)** Copy number (diploid) at the α duplication, based on normalized mean read depth across α, for the All of Us samples split by inversion genotype. Subhaplogroup assignments for H1/H2 samples are shown below; schematics show copy number variation at β and α in H1 and H2 haplotypes respectively. Percentage of H1/H2 samples carrying each subhaplogroup are shown. **C)** H1-H2 recombinant haplotypes identified in the All of Us dataset; 58 haplotypes have an H2 segment on an H1 background and 15 haplotypes have an H1 segment on an H2 background. Stars denote the number of All of Us samples carrying that recombinant haplotype. H1 shown in red/pink; H2 shown in blue/light blue. **D)** H1-H2 recombinant haplotypes (*n*=26) involving combinations of H1 and H2 SDs. Purple regions highlight portions of β and α duplications for which both H1 and H2 copies are present in the recombinant haplotype. The top 3 haplotypes involve a switch from H1 to H2 and likely back to H1 in the SD region. The middle group includes 19 haplotypes with a switch from H2 to H1, with the H2 portion likely carrying an extra H2 α duplication. The bottom 4 haplotypes involve H2 partial (β_p_, α_p_) or full β and α duplications on an H1 background. **E)** Schematic of how β(H2)β(H1) and α(H2)β(H1) haplotypes from (D) may have arisen via H1-H2 crossover events within the single-copy region of the inversion and between SDs. Note that the schematics show H2 haplotypes in reverse orientation, such that the inversion region aligns between H1 and H2 haplotypes.

We found that 7,215 (1.7% of) All of Us individuals carried an H1-H2 recombinant haplotype, defined as a >5-kb stretch of H2 sequence on an H1 background (or H1 sequence on an H2 background). Grouping these recombinant haplotypes by shared switch breakpoints and direction of exchange (H1 into H2 or H2 into H1) yielded 99 distinct recombinant haplotypes (Figure 3C-D). These haplotypes tile the inversion, with each basepair across the inversion region contained in at least one recombinant tract (Figure 3C-D). Four of these recombinant haplotypes are each found in >500 samples (accounting for a total of 5,347 of the 7,215 recombinant samples), primarily in samples with African ancestry (Supplemental Table 2). The remaining recombinant haplotypes are rare (<500 individuals), with 3 recombinant haplotypes occurring in 101-500 individuals and 92 recombinant haplotypes occurring in <100 individuals (Figure 3C-D). We found that most recombinant haplotypes had two switch breakpoints within the single copy region of the inversion (*n*=58 on H1 background, *n*=15 on H2 background), indicative of double crossover or gene conversion events, ranging from 6- to 247-kb in length (Figure 3C). These segments occur in both directions (H2 into H1 and H1 into H2) and are found on all major subhaplogroups (H1.β1, H1.β2, H2.α1, H2.α2) (Figure 3C, Supplemental Table 2). Furthermore, the lengths of these segments in All of Us individuals are consistent with the two-switch tract lengths observed in the sperm data (Figure 2D).

We also identified 26 distinct recombinant haplotypes involving the α and β duplication regions (Figure 3D). For example, two recombinant haplotypes involve H2 sequence that spans the entire β region on an H1 background, resulting in haplotypes with one copy of β (from H2) and one copy of β (from H1), which we denote β(H2)β(H1) (Figure 3D). These haplotypes likely arose via double crossover events, with one crossover occurring between H1 and H2 haplotypes in the single-copy region of the inversion, and the second crossover occurring between H1 and H2 SDs, resulting in haplotypes with β duplications from both H1 and H2 (Figure 3E). We also found recombinant haplotypes that involve an H2 α duplication on an H1 background, resulting in haplotypes with one copy of β (from H1) and one copy of α (from H2), which we denote α(H2)β(H1) (Figure 3D). These α(H2)β(H1) haplotypes are found in >500 individuals (>0.06% frequency) (Figure 3D). The α(H2)β(H1) haplotypes likely arose via double crossover events, for instance, with one crossover occurring in the single-copy inversion region and the second event likely occurring between shared H1 and H2 SDs (Figure 3E). Note that for these double crossover events, homology between H1 and H2 in the inversion region can be obtained via an inversion loop; since single crossovers within an inversion loop are typically inviable, the SDs likely play a critical role in facilitating a second crossover returning to the original haplotype without major chromosomal loss or gain (Figure 3E). Together, these findings indicate that haplotypes with mixed H1 and H2 SDs are not extremely rare, sometimes segregating at appreciable frequencies.

To further explore recombinant haplotypes involving SDs, we analyzed haplotype-resolved genome assemblies from HPRC and HGSVC3. We identified one recombination event between H1 and H2 SDs in an HPRC2 genome^42^. This haplotype exhibits a single crossover event between SDs at the left inversion breakpoint, resulting in a unique SD architecture in which a predominantly H1 haplotype is flanked by γ duplications on both sides (Supplemental Figure 5). This crossover event likely occurred as NAHR between co-linear H1 and H2 SDs (Supplemental Figure 5). Together, these findings of putative crossovers between H1 and H2 SDs in human genomes comport with the H1-H2 crossover events in the sperm data that localize to the SD regions, further underscoring how shared H1 and H2 SDs at breakpoints promote H1-H2 recombination and drive haplotype diversity at this locus.

### Global recombination rate effects of H2

Distinct 17q21.31 haplotypes enabled us to explore the inversion’s phenotypic effects. The H2 inversion orientation is associated with increased maternal global recombination rate, an important fertility-related trait^29^. To evaluate the contribution of the inversion structure to recombination phenotypes, we turned to a previous dataset of 139,416 embryo biopsies with paired parental genotype data^43^. Variants across 17q21.31 are strongly associated with maternal crossover rate (Supplemental Figure 6) and are kept in high LD by the inversion. Using the H2-tagging variant chr17:45900675:C:T, which displays significant association with maternal crossover rate (Supplemental Figure 6), we found that H2 has additive effects on genome-wide crossover rates (Figure 4A). To identify whether this recombination rate effect is global or driven by a subset of chromosomes, we tested for association separately on the chromosome-specific crossover rates. The effect-size estimate for crossover rates when dropping each chromosome remains within a standard error of the effect-size when including all chromosomes (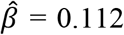, se=0.015; Figure 4A), suggesting that the inversion’s maternal crossover rate effect is not driven by a specific chromosome (*e*.*g*., only driven by chromosome 17). Instead, we found that H2 additively increases crossover rates on each chromosome (Figure 4A).

**Figure 4.**
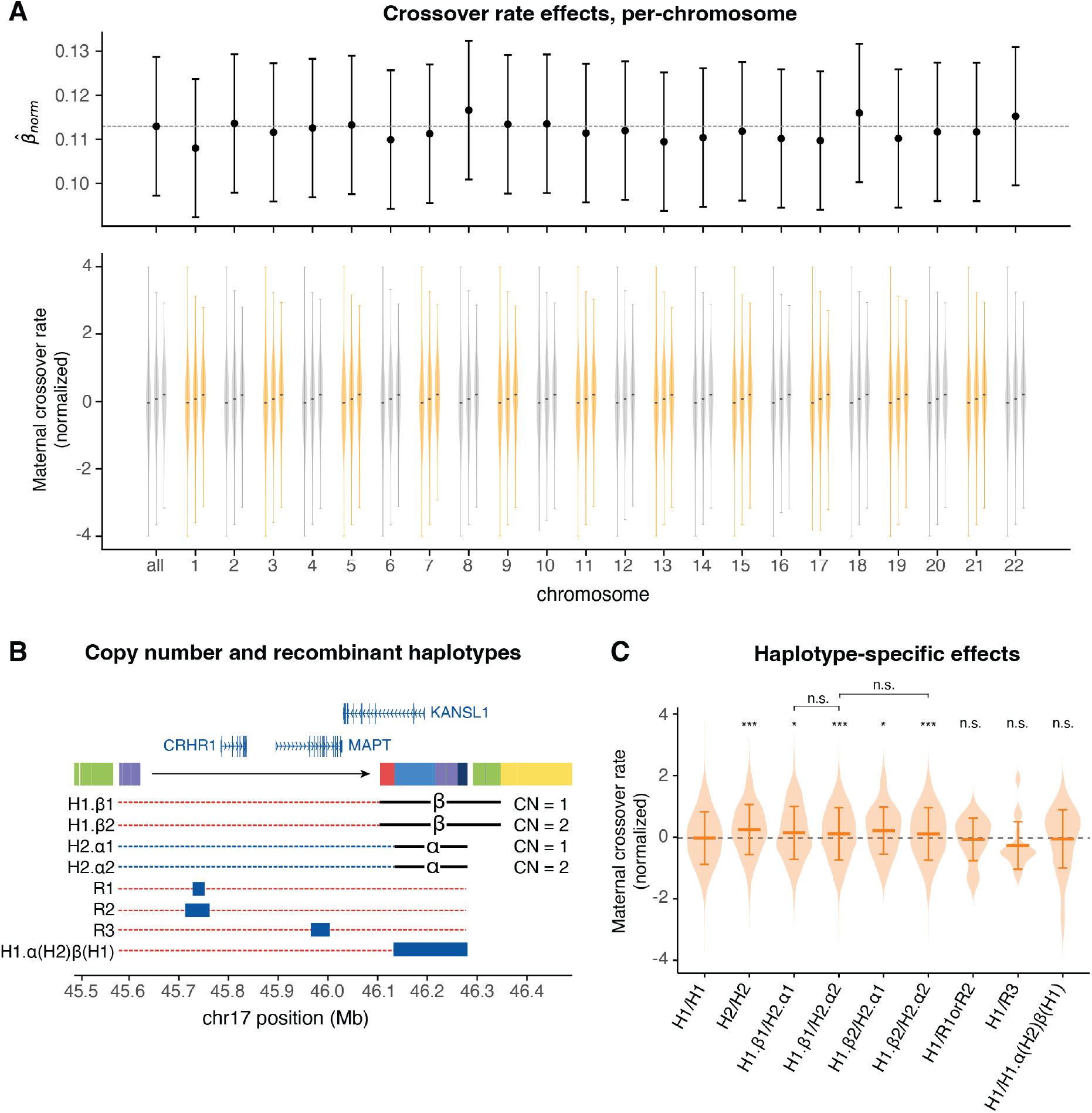
Maternal crossover rate effects of the inversion. **A)** Chromosome-level association of chr17:45900675:C:T with maternal recombination. Top panel reflects effect sizes from linear regression (including age and 20 genetic principal components), leaving one chromosome out. Bottom panel shows phenotype distribution as a function of genotypes (CC, CT, TT), where C tags H1 and T tags H2. **B)** Distinct 17q21.31 haplotypes for dissecting the inversion’s crossover rate effects include copy number (CN) variants for β and α duplications and H1-H2 recombinant haplotypes. Haplotype background is indicated by dashed lines, with H1=red, H2=blue. **C)** Comparison of maternal recombination rate stratified by inversion subhaplogroup. The dashed line indicates the mean recombination rate of the H1/H1 individuals for comparison as the null baseline. Significance levels (t-test): n.s. = not significant, * = p < 0.05; *** = p < 0.001, all subhaplogroups compared to H1/H1 baseline (Supplemental Table 3), and CN 1 v. 2 for β and α duplications.

We next tested whether 17q21.31 breakpoint SDs contribute to the inversion’s effects on crossover rates. Since inversion status (H1 v. H2) is highly correlated with *KANSL1* copy number (Figure 3B), the inversion’s maternal crossover rate effect may be driven by duplication status rather than inversion status. We assigned copy number for α and β duplications in H1/H2 parents using allelic balance at informative SNPs (Supplemental Figure 7) and compared maternal crossover rates across subhaplogroup assignments (Figure 4B,C). There were no differences in maternal crossover rates by α copy number on H2 (H1.β1/H2.α1 v. H1.β1/H2.α2: *p* = 0.257) or β copy number on H1 (H1.β1/H2.α2 v H1.β2/H2.α2: *p* = 0.330) (Figure 4C), suggesting that inversion status, and not duplication status, drives maternal crossover effects.

Fine-mapping within an inversion is impeded by the strong LD it causes. We took advantage of H1-H2 recombinant haplotypes, which break apart LD at the 17q21.31 locus, to map the recombination rate effects within the inversion. We identified 17 maternal samples carrying an H1.α(H2) β(H1) haplotype, in which the α region from H2 has recombined onto an H1 background (Figure 4B). This haplotype shows no increase in crossover rates compared to H1/H1 (*p* = 0.961) (Figure 4C), suggesting that H2 variants in the α region of *KANSL1* are unlikely to be causal of the crossover effect (but noting that we have limited power to detect this haplotype’s effect). We also identified 9 maternal samples carrying recombinant segment 1 (R1), 4 carrying recombinant segment 2 (R2) and 17 carrying recombinant segment 3 (R3) respectively, in which H2 sequence has recombined onto an H1 background (Figure 4B). We did not find differences in crossover rates for individuals carrying each recombinant segment (R1 and R2; R3) compared to H1/H1 individuals (noting the limited power to detect an effect) (Figure 4C). These recombinant haplotypes therefore help refine the causal region for H2’s effects on maternal crossover rates, but larger sample sizes are required to distinguish between recombinant haplotype effects.

### A history of genetic exchange between H1 and H2 haplotypes

We next explored the evolutionary history of the inversion using ancestral recombination graphs (ARGs). ARGs capture the genealogical relationships among haplotypes at each segment of the genome, enabling detailed analyses of 17q21.31 haplotypes across the inversion and through time. We used the Human Genome Diversity Project (HGDP) and 1KGP datasets, which include H2 haplotypes from across continental groups and multiple recombinant haplotypes (Supplemental Figure 8). We focused evolutionary analyses on four of the most common recombinant haplotypes (R1-R4) with segments spanning 23-49 kb (Figure 5A, Supplemental Table 2). We also included a putative 30-kb recombination event at the *CRHR1* gene, a region which shows striking depletion for H1-H2 differentiation (Figure 1C) suggestive of a fixed recombinant haplotype^30,33^. Using a panel of ~150 haplotypes chosen to capture the diversity at the 17q21.31 locus and include R1-R4 recombinant haplotypes, we inferred ARGs across the inversion region with SINGER^44^ and estimated time to most recent common ancestor (TMRCA) between haplotypes of interest. We note that our TMRCA estimates are robust to varying input parameters and haplotype panels (Supplemental Figure 9). By excluding recombinant haplotypes and the *CRHR1* 30-kb region, we estimated that H1 and H2 diverged ~4 Mya (95% credible interval: 3.85-4.15Mya), H2 haplotypes across modern humans coalesced ~250 kya (95% credible interval: 221-290 kya) and H1 haplotypes coalesced ~1.4 Mya (95% credible interval: 1.28-1.53 Mya) (Figure 5B).

H1 and H2 haplotypes consistently form distinct clades across the inversion region except at the *CRHR1* 30-kb segment (Figure 5C, Supplemental Figure 10). In this *CRHR1* region, H2 haplotypes form a clade nested within H1 haplotypes (Figure 5C), suggesting that an H1 segment recombined into an H2 haplotype and subsequently fixed within all H2 haplotypes. With the ARGs, we estimated the TMRCA between H2 and H1 haplotypes in this region, inferring the timing of this *CRHR1* recombination event at ~520 kya (95% credible interval: 327-836 kya) (Figure 5B), occurring before the spread of H2 haplotypes within modern humans. Lineages tracked within the inferred ARGs also reveal the evolutionary history of R1-R4 events. Trees from within the R1 segment revealed that R1 haplotypes form a distinct clade from R2 haplotypes (Figure 5C), suggesting that R1 and R2 were independent events involving an H2 segment recombining onto an H1 background in the same part of the inversion (Supplemental Figure 10). R1 haplotypes form a sister clade to H2 haplotypes (Figure 5C), with an estimated divergence time from H2 of ~330 kya (95% credible interval: 192-564 kya) (Figure 5B); R2 haplotypes form a clade nested within H2 haplotypes (Figure 5C), with an estimated divergence time to nearest H2 haplotype of ~100 kya (95% credible interval: 75-150 kya) (Figure 5B). In contrast, R3 and R4 appeared as a single sister clade to H2 (Figure 5C), suggesting that one H2 into H1 recombination event led to R3 occurring ~450 kya (95% credible interval: 325-640 kya) (Figure 5B, Supplemental Figure 10), and then a subsequent recombination event between R3 and H1 haplotypes led to the smaller R4 tract. Taken together, these results suggest that gene flux between H1 and H2 has occurred over at least the last 500k years, with genetic exchange occurring from H1 into H2 and also H2 into H1 (Figure 5D).

**Figure 5.**
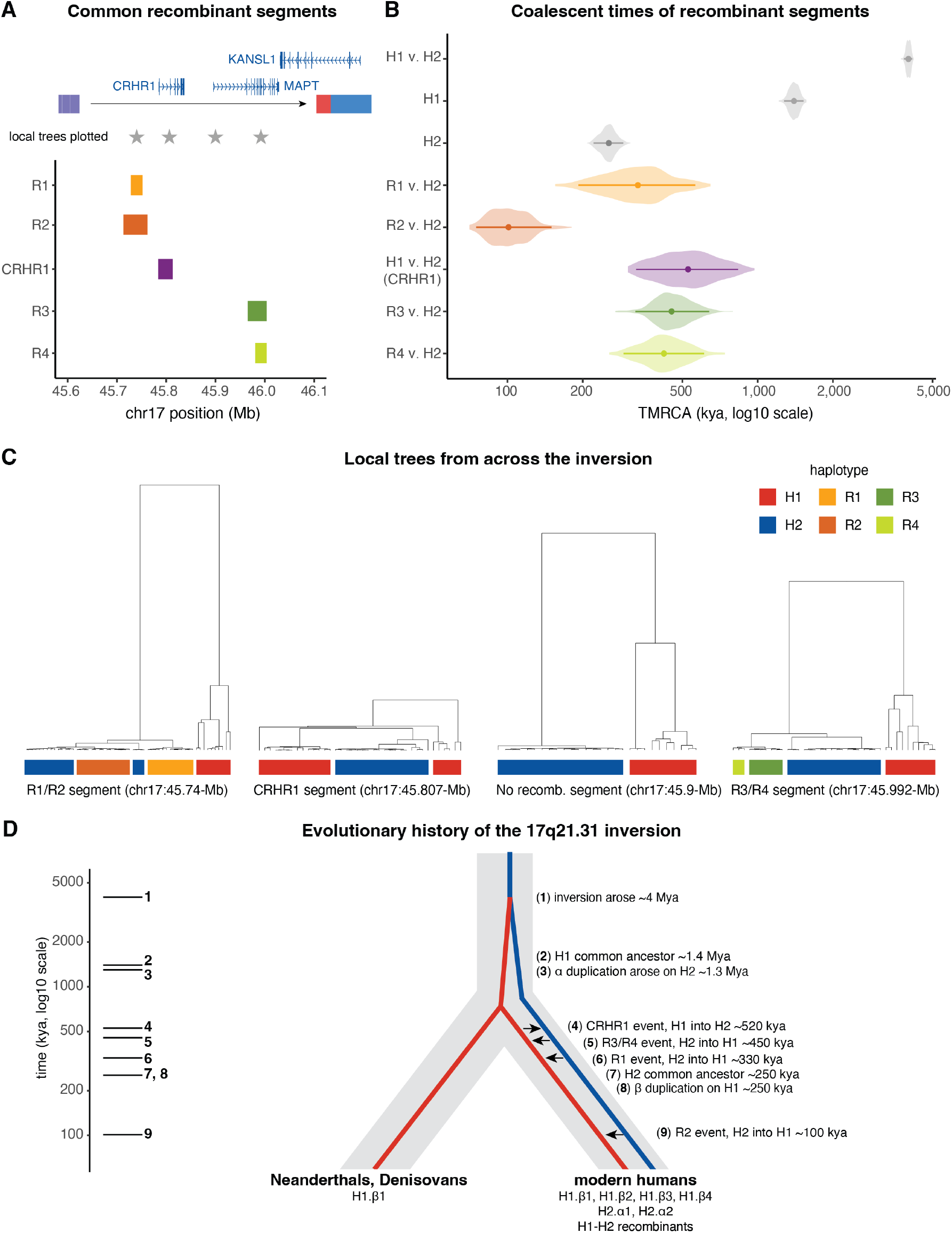
Evolutionary history of the inversion and its gene flux events. **A)** Location of 4 common H1-H2 recombination events (R1-R4) and a fixed recombination event at the *CRHR1* gene. Schematic of the inversion’s single copy region (black arrow) and flanking SDs is shown above, with its three main genes. The locations of four local trees (plotted in C) are shown with gray stars. **B)** Coalescent time estimates from ARGs for H1 v. H2 haplotypes, H1 haplotypes, H2 haplotypes, R1-R4 v. H2 haplotypes within each recombinant segment, and H1 v. H2 within the *CRHR1* recombination region. Posterior median is shown as dot, with 95% credible interval shown as lines and violin plots showing posterior distribution from the full set of ARGs. **C)** Local trees sampled from the ARGs for four locations across the inversion region. Tips are highlighted by haplotype group to show relationships amongst H1, H2 and recombinant haplotypes. **D)** Diagram of the evolutionary history of the inversion, with major events listed, including α and β duplication time estimates from ^30^ and timing of inversion origin, spread and H1-H2 recombination events estimated in this study.

## Discussion

Our study provides a comprehensive analysis of the 17q21.31 inversion’s effects on recombination, demonstrating that this human inversion is an incomplete barrier to local recombination. Instead, two-switch events, as well as breakpoint-mediated exchange, drive ongoing gene flux between H1 and H2 haplotypes. Unlike single crossovers within inversions, which lead to unbalanced gametes, two-switch events allow for gene flux without sequence loss^45^ (Supplemental Figure 11). Two-switch events between H1 and H2 measured in single sperm genomes occurred at an unexpectedly high frequency, in >1 out of 1,000 meioses. This rate exceeds published genome-wide estimates of long gene-conversion tracts of comparable size^39,40^, but it includes both gene conversion and double crossovers. However, we hypothesize that gene conversion is the main contributor: recent studies suggest a long (multi-kilobase) component to gene conversion in addition to its typical short <100 bp tract length^39,46^ and we estimated that the probability of gene conversion is much higher than double crossovers for the observed tract lengths. In *Drosophila*, gene conversion events occur between inversion haplotypes at rates comparable to collinear regions^24,47,48^, but they are much shorter tracts (< 500 bp) compared to what we observed at 17q21.31. Thus, our results raise the question of whether kilobase-scale gene flux is a common feature of large inversion polymorphisms.

Our findings demonstrate how breakpoint architectures provide a distinct route for gene flux at inversions. 17q21.31 is characterized by complex SD architectures at inversion breakpoints, including co-linear SDs found between H1 and H2 haplotypes within the left breakpoint region and within the right breakpoint region. We found that these co-linear SDs shared between H1 and H2 haplotypes promote single crossovers via NAHR, also occurring in >1 out of 1,000 meioses. Remarkably, these single crossovers do not yield major chromosomal loss or gain because they occur as NAHR within left (or right) breakpoint regions of both H1 and H2, rather than homologous recombination via an inversion loop (Supplemental Figure 11). We found that NAHR between these SDs can generate novel locations and copy number of SDs, likely explaining the extensive variability of 17q21.31 breakpoint architectures^30,32,33^. However, these SDs also make the region susceptible to structural rearrangements: NAHR has been shown to occur between left and right breakpoint SDs, and results in a ~500-kb deletion at 17q21.31^35^. This microdeletion causes Koolen-de Vries syndrome – a developmental disorder that occurs in 1 out of 16,000 individuals^49^ or ~1 out of 3,000 H2 carriers^50^. In addition, 17q21.31 SDs likely promote the inversion’s recurrence and mutability across primates via NAHR^33,34^. Therefore, the same breakpoint SDs that contribute to inversion formation and genomic instability also facilitate genetic exchange between arrangements, suggesting a more central role of breakpoint architectures in inversion evolution.

Even occasional gene flux can profoundly alter the long-term evolution of an inversion. We found that ongoing gene flux has generated abundant haplotype diversity, with at least 99 distinct H1-H2 recombinant haplotypes segregating in the human population, a finding that substantially expands upon previously identified recombinant haplotypes at this locus^33,51^. These recombinant haplotypes provide a unique lens into the inversion’s evolutionary history; by inferring coalescence times of recombinant haplotypes with ancestral recombination graphs, we estimated that H1 and H2 were co-segregating ~500kya, before the out-of-Africa migration. This pushes the age of H1 and H2 co-segregation far deeper than previous estimates of ~136kya based on H2 coalescence times^30^. It is possible that the two haplotypes have co-existed for even longer (e.g., since the inversion arose approximately 2-4Mya), but different demographic models, such as deep population structure or ghost introgression^52–54^, might explain the high divergence between H1 and H2. Furthermore, the coalescence times of recombinant haplotypes indicate that gene flux has been ongoing for the last 500k years. One possible consequence of this gene flux is a reduction in the inversion’s fitness costs. Inversion polymorphisms are predicted to accumulate deleterious mutational loads over time, since purifying selection is less effective without recombination, but gene flux can reduce mutational loads through breaking apart LD at an inversion^26^. Therefore, it is possible that gene flux helped maintain the 17q21.31 inversion over the last half a million years. For example, the highest frequency recombinant event is a 30-kb H1 tract that is fixed on the H2 background, spanning the introns and exons of *CRHR1*, a gene involved in the stress response^31^. We estimated that this event occurred ~520 kya, before the coalescence of H2 ~250 kya in modern humans, perhaps facilitating the spread of H2 by removing deleterious mutations accumulated at *CRHR1*. That inversions can be permissive to gene flux may allow them to confer the benefits of suppressed recombination (e.g., through linking co-adaptive alleles) without severe fitness costs, motivating measurements of gene flux at inversion polymorphisms found within a broad range of species.

The natural recombinant haplotypes generated by gene flux facilitate dissection of the phenotypic effects of inversions, which are typically obscured by high LD. Through characterizing distinct 17q21.31 haplotypes, we helped resolve the inversion’s effects on maternal crossover rates, an important fertility-related phenotype. Our analysis of a large parent-embryo dataset^43^ revealed that H2 has a *trans* effect on maternal crossovers, increasing crossover rates on each chromosome (including chromosome 17 where it resides). In *Drosophila*, an interchromosomal recombination effect of inversions has also been observed^55^ and is attributed to structural features of inversions: an inversion can delay meiotic checkpoints due to lack of homologous alignment or crossovers, leading to a longer period for crossovers to occur during meiosis^47,56^. We demonstrate that structural features are unlikely to be causal of 17q21.31’s global recombination effects. First, H2 additively increases maternal crossover rates, such that the highest recombination rates are found in H2 homozygotes, a point of evidence against the delayed meiotic checkpoint hypothesis. Second, we show that *KANSL1* duplications on H1 and H2 are not associated with increased crossover rates. A recent study showed that *KANSL1* duplications have rapidly increased in frequency on both H1 and H2 haplotypes over the last 10k years, suggesting that these duplications may be targets of selection^33^; however, our findings argue against the role of breakpoint *KANSL1* duplications in driving this fertility-related phenotype. Rather than structural features, H2 may impact maternal crossover rates via mutation(s) harbored within the inversion, which we began to explore through comparisons across recombinant haplotypes, but the low frequency of recombinant haplotypes limited our power to fine-map. Nevertheless, this approach provides a compelling framework for resolving inversion phenotypes, and motivates large-scale, targeted association studies with recombinant haplotypes.

In summary, the 17q21.31 inversion is a fascinating locus, as it is one of the human genome’s most dynamic and ancient polymorphisms and it has profound effects on both local and global recombination rates. Our study reveals how gene flux, driven in part by its complex breakpoints, has occurred over at least half a million years and shaped the diversity and evolution of the 17q21.31 inversion, providing a clue into the puzzling persistence of inversion polymorphisms.

## Methods

### Segmental duplication annotations

Segmental duplications (SDs) were localized in the T2T-CHM13 genome using the UCSC genome browser segmental duplication track and read depth signal from short-read sequencing (see below). Duplication names correspond to those previously described^32^. A duplicon library fasta covering these regions was built by extracting the SD regions from T2T-CHM13 with *samtools*. Haplotype-resolved genome assemblies were downloaded from HPRC2^36,42^ and HGSVC3^37^. The 17q21.31 region was extracted from each assembly by aligning the assembly to T2T-CHM13 sequences upstream (chr17:44.9-45Mb) and downstream (chr17:48.7-48.8Mb) of the inversion using *minimap2* (-x asm20). SDs were annotated by querying the assemblies against the duplicon library fasta with *blastn*, keeping hits with >90% identity and >500-bp alignment length. The 17q21.31 region was also extracted from GRCh38 and annotated for the SDs with *blastn*. To visualize the inversion, a representative H2 haplotype from HPRC was aligned to T2T-CHM13 with *minimap2* (-x asm5) and then plotted with SVbyEye ^57^, retaining alignments >200-kb.

### Identifying H1-H2 tag SNPs

To identify SNPs tagging the H1 versus H2 haplotypes, we first genotyped 1KGP genomes for the inversion using PCA. Biallelic SNPs at the 17q21.31 locus were extracted from the 1KGP high-coverage phased vcf (GRCh38) with *bcftools* and loaded in *scikit-allel*. Local PCA was performed from chr17:45-47Mb (GRCh38) using 50-kb windows with *allel*.*pca* (scaler=patterson), with sign of PC1 aligned to prior window to minimize axis flips. Overall PCA was performed on SNPs from chr17:45.5-46.25Mb (GRCh38) and samples were assigned to inversion genotype based on PC1 score. Tag SNPs were identified as biallelic sites with H1/H1 allele frequency < 5% and H2/H2 allele frequency > 95% across all 1KGP samples, yielding a total of 2,461 SNPs across the 17q21.31 locus.

### Recombination rates from single-cell sperm-sequencing

We analyzed single-cell sperm DNA sequencing from 20 donors using data from Bell *et al*. 2020^38^ (accessed from dbGaP), which includes low-coverage (0.01x) whole-genome sequencing from 974 - 2,274 cells per donor. Sequencing reads from per donor fastq files were aligned to GRCh38 using *bwa mem*. We first inferred donor inversion haplotypes. To do so, we created donor-level vcfs using GATK *HaplotypeCaller* (per donor), *CombineGVCFs* and *GenotypeGVCFs*. We selected biallelic SNPs and performed hard-filtering based on GATK recommendations (QD < 2, FS > 60, MQ < 40, MQRankSum < −12.5, ReadPosRankSum < −8, SOR > 3). H1 versus H2 inversion haplotypes were assigned to each donor based on donor genotypes at tag SNPs, noting that all donors had consistent genotypes for >98% of the tag SNPs. 15 donors were assigned as H1/H1 and 5 donors as H1/H2. For the H1/H2 donors, we assigned subhaplogroups based on copy number at α, β and γ regions. We note that the full β duplication (GRCh38 chr17:46.106-46.281 Mb) contains α (chr17:46.135-46.219 Mb), so for copy number analyses of β, we used the unique portion of the β duplication (chr17:46.106-46.135 Mb), which we hereafter refer to as the β’ region. To infer copy number, we calculated read depth in 1-kb windows across the locus using *samtools depth* from the per donor bam files; we normalized read depth by average depth from chr17:44.0-44.2Mb, a single copy genomic region outside of the inversion. We also used mean allelic balance (defined as depth at H2 allele/total depth) for tag SNPs in α and β’ regions to infer number of H1 v. H2 α and β’ duplicates. We obtained diploid copy number for γ by summing mean depth at both γ copies in GRCh38; for diploid γ copy number of 3, we inferred the most likely assignment as one H1 γ, two H2 γ based on the low frequency of H2.γ1 subhaplogroups^32,33^.

We identified recombination events on chr17 in single sperm cells using *rhapsodi*, a method for crossover detection from low-coverage single-cell gamete sequencing^58^. To create *rhapsodi* input files, we first selected heterozygous SNPs from each donor using GATK SelectVariants, excluding SNPs in low mappability and SD regions from Genome in a Bottle. We then genotyped single sperm cells using reads in the BAM files that overlapped donor-specific heterozygous SNPs, with read barcode informing cell identity, to create input genotype matrices. We excluded low-quality sperm cells or those with chr17 aneuploidy calls determined from^38^. We ran *rhapsodi* for chr17 with *avg_recomb* as 1.5 and *coverage* as 0.01, masking the full α, β, γ segmental duplication region (chr17:46.088-46.707Mb) to avoid spurious recombination calls. *Rhapsodi* uses the single-cell sperm genotypes to phase donor SNPs and then estimate crossover breakpoints in each sperm cell, returning the start and end position for estimated crossover events.

Sperm cells with single-crossover events within the inversion region were identified from the *rhapsodi* crossover calls. We selected sperm cells with only one crossover event contained within 500-kb of the inversion (chr17:44.915 - 47.207Mb), excluding cells with more than one crossover in the extended region (chr17:43.915 – 48.207Mb). To estimate recombination rates at the inversion locus, we calculated per-donor inversion region recombination rate as the number of cells with a crossover event in the region divided by the number of cells in which a crossover event was detectable (i.e., cells with genotypes for ≥2 informative SNPs, separated by ≥500-kb, within the region). Recombination rates were compared between H1/H1 and H1/H2 genotypes using a permutation test, in which genotype was shuffled across donors (*n*=10,000), weighting by the total number of crossover-detectable cells per genotype. We used genotype at tag SNPs to assign H1 v. H2 haplotypes to sperm cells with single crossover events; for cells with single crossovers on an H2 background, we flipped the coordinates of SNPs within the inversion region to resolve and plot the crossover locations. To visualize crossovers occurring within SD regions, we projected crossover confidence intervals onto HPRC assemblies representative of the sperm donors’ 17q21.31 subhaplogroups. For example, a sperm donor with subhaplogroup H2.α2.γ2 was visualized with HPRC haplotype-resolved assembly harboring H2.α2.γ2. To liftover GRCh38 coordinates for each crossover event into each haplotype-resolved assembly’s coordinate space, we used the shared SD boundaries as anchors.

Two-switch events were also detected using the *rhapsodi* crossover calls. The high SNP density between H1 and H2 haplotypes facilitated two-switch event detection in H1/H2 donors only (since informative SNP density was insufficient in H1/H1 donors). For these events, we required a sperm cell to have two crossover events within the inversion region (chr17:44.915 - 47.207Mb), with at least 3 independent SNPs (located >200-bp apart) supporting the tract. We visualized two-switch events using the confidence intervals for both crossover events, denoting likely crossover locations.

### Inversion genotypes and copy number variation in the All of Us dataset

We genotyped the 414,830 All of Us participants for the inversion using the H1-H2 tag SNPs. Analysis of the All of Us dataset was performed on the Researcher Workbench. We loaded the Hail matrix table (ACAF, v8), which contains data for variants with >1% frequency in a population or with allele count >100 from the whole-genome sequences, and performed per sample counts of genotypes at 2,286 tag SNPs in Hail. A sample was assigned to an inversion genotype (H1/H1, H1/H2, H2/H2) if ≥98% of its tag SNPs supported one inversion genotype.

We performed copy number analysis of the α and β’ duplications for the All of Us samples. We used the VariantDataset (VDS, v8), which contains local allele depth (LAD) at each sample’s variant sites (0/1 or 1/1 genotypes). We calculated per sample mean depth across variant sites for α, β’, and a normalization window that does not contain duplications (chr17:44.0-44.2 Mb). Diploid copy number for α (or β’) was estimated as two times mean depth at α (or β’) divided by mean depth at the normalization window.

For H1/H2 samples, we also assigned haploid copy number of α and β’ duplications by analyzing allelic balance. We first identified SNPs within α and β’ regions that were diagnostic of H1 v. H2 haplotypes; we filtered the Hail matrix table for biallelic SNPs with >90% frequency in H1/H1 samples and <10% frequency in H2/H2 samples, or vice versa, yielding 147 diagnostic SNPs for α, and 131 diagnostic SNPs for β’. We then computed per sample allelic balance, defined as depth at the H2 allele divided by total depth at the site, at the diagnostic SNPs and took the mean allelic balance at α SNPs and β’ SNPs. The H1/H2 samples were then clustered based on normalized depth at α and β’ and mean allelic balance at α and β’; samples were assigned to subhaplogroup based on nearest centroid, defined from known subhaplogroups, in scaled Euclidean space (Supplemental Figure 4). For example, we expect H1.β1/H2.α2 samples to have diploid copy number of 2 for β’ and 3 for α, and allelic balance of 0.5 for β’ and 0.66 for α.

### Recombinant haplotypes in the All of Us dataset

We identified H1-H2 recombinant haplotypes in the All of Us dataset based on genotypes at the H1-H2 tag SNPs. Using the Hail matrix table, we first identified samples with segments of ≥30 contiguous tag SNPs that differ from their mode tag SNP genotype, allowing for up to a 5 tag SNP gap within the segment. We ensured that these haplotypes represented H1-H2 recombination, and not deletions, by checking allelic depth across the haplotype: we excluded 6 samples whose haplotypes appeared to be deletions, with mean depth in the putatively recombined segment <70% of mean depth in the rest of the tag SNP region. From the remaining 7,215 recombinant samples, we identified a set of unique recombinant haplotypes by merging recombinant haplotypes with shared H1-H2 switch breakpoints (within 5-kb of each other), resulting in 99 distinct recombinant haplotypes. We then inferred haploid copy number and identity (H1 v. H2) of α and β’ duplications in the recombinant haplotypes based on diploid α, β’ copy number and mean allelic balance (if heterozygous) in samples carrying that recombinant haplotype. We recorded ancestry groups for each recombinant haplotype based on All of Us ancestry predictions.

### Maternal crossover rate association analysis

To analyze the inversion’s association with maternal crossover rates, we used a previous dataset of 139,416 embryo biopsies with paired parental genotype data (Carioscia *et al*. 2026)^43^, restricting to maternal individuals with >3 euploid embryos (*n* = 15,333). The lead variant for maternal recombination rate in this region is chr17:46287153:C:G, but for most of our analyses, we consider the H2-tagging variant chr17:45900675:C:T, which also displays significant association with maternal recombination rate (Supplemental Figure 6). To test whether the crossover rate effect of the inversion is unevenly distributed across chromosomes, we performed association analysis separately on the chromosome-specific crossover rate. We specifically transformed the crossover counts using inverse-rank normal transformation so all effect-sizes can be interpreted in units of standard deviation from the mean recombination rate. We tested for this effect by excluding a focal chromosome from crossover counts and using linear regression with the genotype as the primary predictor and age and twenty genetic ancestry principal components as covariates.

We next assigned copy number at α and β duplications in H1/H2 individuals. Since individuals in this dataset were genotyped with microarrays, we inferred allelic balance at 4 α SNPs (rs2732589, rs1918792, rs2732631, rs2668692) and 2 β’ SNPs (rs17660907, rs4383188) using raw array probe intensity values. At these SNPs, we computed the ratio of probe intensities, B/(A+B) where B is the intensity of the H1 allele and A is the intensity of the H2 allele; for one SNP from each of α and β’ SNP sets, the H1 allele corresponded to probe A rather than probe B, so we instead computed A/(A+B). Taking the average ratio of probe intensities across α SNPs and β’ SNPs thus cancelled out technical variation in A/B channel intensity. Using the average ratio of intensities as a proxy for allelic balance at α and β’, we inferred number of H1 v. H2 copies of α and β in H1/H2 individuals (Supplemental Figure 7). Using these assignments, we compared the maternal recombination rate across individuals, stratifying by subhaplogroup (Supplemental Table 3).

We also identified maternal individuals in this dataset with H1-H2 recombinant haplotypes. In analyzing allelic balance for α and β’ SNPs in H1/H1 individuals, we found 17 individuals with allelic balance of 1 for β’ SNPs and 0.66 for α SNPs, indicating that these individuals carry an H1 haplotype with 1 H1 β copy and 1 H2 α copy (which we denote H1.α(H2)β(H1)). We also used H1-H2 tag SNPs to identify individuals carrying common recombinant haplotypes (i.e., H2 sequence at segment 1 (chr17:45,728,160-45,751,990); segment 2 (chr17:45,713,030-45,761,890); and segment 3 (chr17:45,965,550-46,003,700) on an H1 background). We then compared maternal crossover rates between each recombinant group and H1/H1 individuals.

### Dating H1-H2 recombination events with Ancestral Recombination Graphs

To characterize the evolutionary history of the H1-H2 recombination events, we estimated coalescence times using ancestral recombination graphs (ARGs). To do so, we first identified recombinant haplotypes in the 1KGP and HGDP datasets (*n*=4,091 samples). We extracted the H1-H2 tag SNPs from a combined vcf of 1KGP and HGDP samples (https://gnomad.broadinstitute.org/downloads#v4-variants) using *bcftools* and then assigned inversion genotype and recombinant haplotypes as described above for the All of Us data. We selected samples for ARG analyses based on inversion genotype and recombinant status: to maximize H2 and recombinant haplotype representation within a feasible sample size, we included all H2 African haplotypes, H1-H2 recombinant haplotypes (up to 3 per haplotype per population), at least 1 H1 haplotype per African population, and additional H1 and H2 haplotypes from different continental groups. In total, we included 153 haplotypes (n=97 H1, 18 H2, 13 R1, 15 R2, 7 R3, 3 R4). We used *bcftools* to extract biallelic SNPs from chr17:45.5-46.1Mb (GRCh38) from the phased 1KGP & HGDP vcf, requiring no missing genotypes. We inferred ARGs for the inversion region (chr17:45.5-46.1 Mb) using SINGER v0.1.8-beta, which samples ARGs from the posterior distribution, providing uncertainty characterization^44^. As input parameters, we used a mutation rate of 1.25×10^−8^ per bp per generation^59^, ratio of recombination to mutation rate of 0.597 (based on mean recombination in the region of 7.46×10^−9^ from the SHAPEIT b38 genetic recombination maps), effective population size (*N*_*e*_) of 10,000, generation time of 28 years, and sites treated as unpolarized (polar=0.5). We ran 4 independent chains, drawing 100 posterior ARGs per chain with thin=100 and discarding the first 30 samples of each chain as burn-in.

We estimated coalescence times for each event from the ARGs, using trees from the region of interest. Specifically, we computed the mean TMRCA from each posterior ARG using *n* random marginal trees spaced across the region of interest (*n* = 600 for full inversion region, *n* = 100 for recombination events). For recombinant segments (including the *CRHR1* region), we excluded trees that had spans intersecting with the first and last 10% of the segment region to ensure that trees reflected relationships within the recombinant segment itself. Using the 280 post burn-in ARGs, we recorded the posterior median divergence time for each event of interest and the 95% credible interval. We estimated the TMRCA of H1 and H2 haplotypes, TMRCA of the largest monophyletic subset of H1 haplotypes, and TMRCA of the largest monophyletic subset of H2 haplotypes in the reference panel using the full inversion region (chr17:45.6-46.07 Mb) but excluding the *CRHR1* recombinant segment (chr17:45.783-45.813 Mb). For recombinant haplotypes, the local trees showed that R1 haplotypes were sister to the H2 clade within the R1 segment, so we estimated the timing of R1 recombination event as the TMRCA of R1 haplotypes to the H2 clade, using trees from within the R1 segment. We also found that R3 and R4 haplotypes were sister to the H2 clade and thus used TMRCA of R3 or R4 haplotypes to H2, using trees from within segments R3 and R4 respectively. In contrast, we found that R2 was nested within the H2 clade, so we estimated the timing of the R2 recombination event as the mean TMRCA of R2 haplotypes to the nearest H2 haplotype, using trees within the R2 segment. Similarly, in the *CRHR1* recombination segment, we found that all H2 haplotypes were nested within H1, so we estimated the timing of this recombination event as the TMRCA of all H2 haplotypes to the nearest H1 clade, requiring at least 3 H1 haplotypes in the nearest clade.

We tested the sensitivity of our coalescence estimates by re-running SINGER with different input parameters and sample sets. We let *N*_*e*_ = 5,000 and 20,000, compared to baseline of 10,000. We let the ratio of recombination to mutation rate = 0.008, 0.08, 0.4, 0.96 (corresponding to recombination rates of 1×10^−10^, 1×10^−9^, 5×10^−9^, 1.2×10^−8^ per basepair per generation, assuming a mutation rate of 1.25×10^−8^), compared to baseline of 0.597 (corresponding to recombination rate of 7.46×10^−9^); these variable recombination rates span different estimates at this region (e.g., 5.8×10^−9^ from ^39^; 3.7×10^−9^ from ^60^). To assess the sensitivity of our estimates to which haplotypes were included in our reference panel, we created alternative panels of equal size (*n*=150) and re-ran ARG inference and coalescence time inference on each panel. For H1-H2, within H1, and within H2 divergence times, we used 3 balanced panels of 75 randomly selected H1 and 75 randomly selected H2 haplotypes and 3 panels of 150 randomly selected haplotypes from all H1 and H2 haplotypes that more closely reflect H1 and H2 frequencies. For recombination event times, we used 3 panels per event of all recombinant haplotypes for that segment plus a random selection of H1 and H2 haplotypes to reach 150 total; in addition, we used 3 panels per event of a random selection of 150 haplotypes, requiring at least 1 recombinant haplotype of the segment of interest to be included. For each condition and coalescence time estimate, we report the posterior median and 95% credible intervals from the SINGER run, which included 280 post-burn-in ARGs (4 independent chains, 100 posterior samples per chain, excluding the first 30 as burn-in) (Supplemental Figure 9).

## Acknowledgements

We thank the NIH All of Us Research Program for making available the sequencing data examined in this study. We acknowledge All of Us participants for their contributions, without whom this research would not have been possible.

## Data availability

Raw sperm sequencing data from Bell *et al*. 2020^38^ can be accessed via dbGaP (study accession number phs001887.v1.p1). The All of Us sequencing data are available from the All of Us Research Program in the Controlled Tier (https://www.researchallofus.org). The 1KGP and HGDP sequences are publicly available from https://www.internationalgenome.org. The HPRC and HGSVC genome assemblies are publicly available from https://humanpangenome.org and https://www.internationalgenome.org/data-portal/data-collection/hgsvc3.

---

**Supplemental Figure 1.**
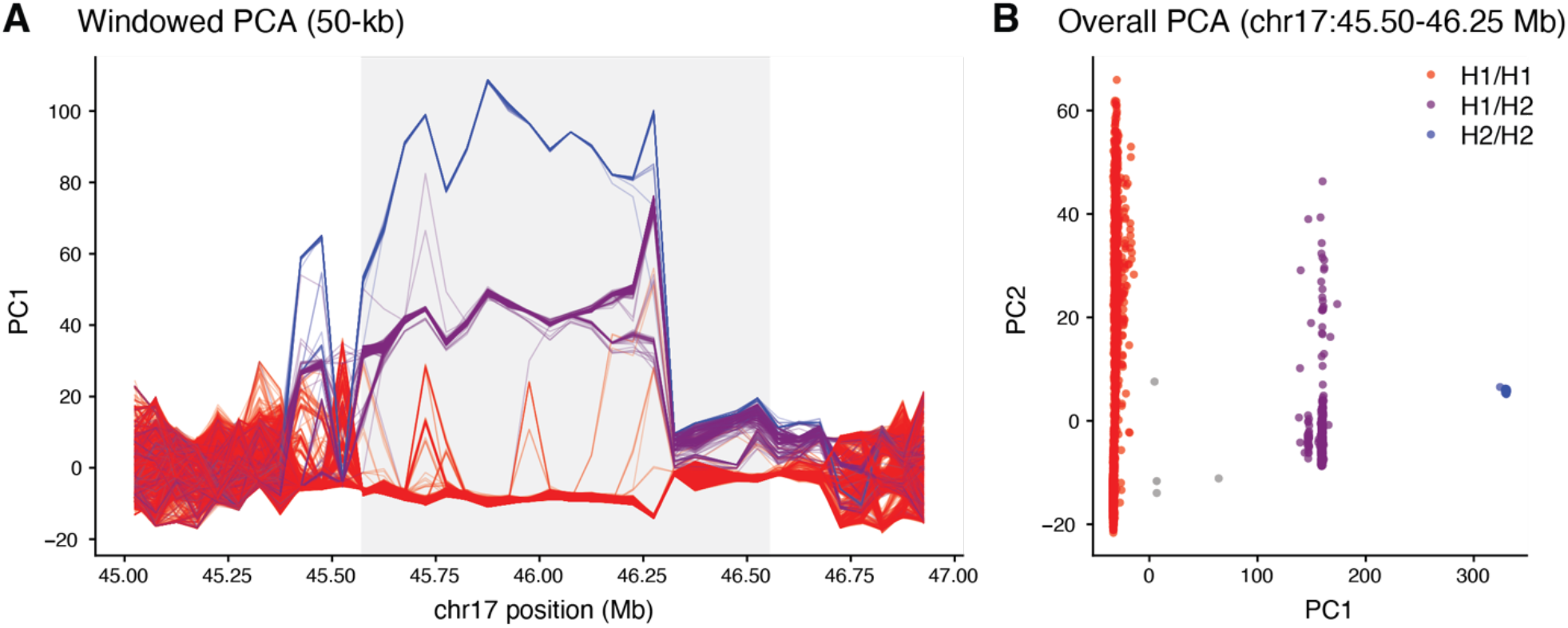
Genotyping the inversion in the 1KGP samples with PCA. **A)** Local PCA using 50-kb windows along GRCh38 for the 3,202 1KGP samples. The inversion region is highlighted in gray. **B)** Overall PCA for the 1KGP samples using SNPs from chr17:45.5-46.25 Mb, with samples colored by inferred inversion genotypes.

**Supplemental Figure 2.**
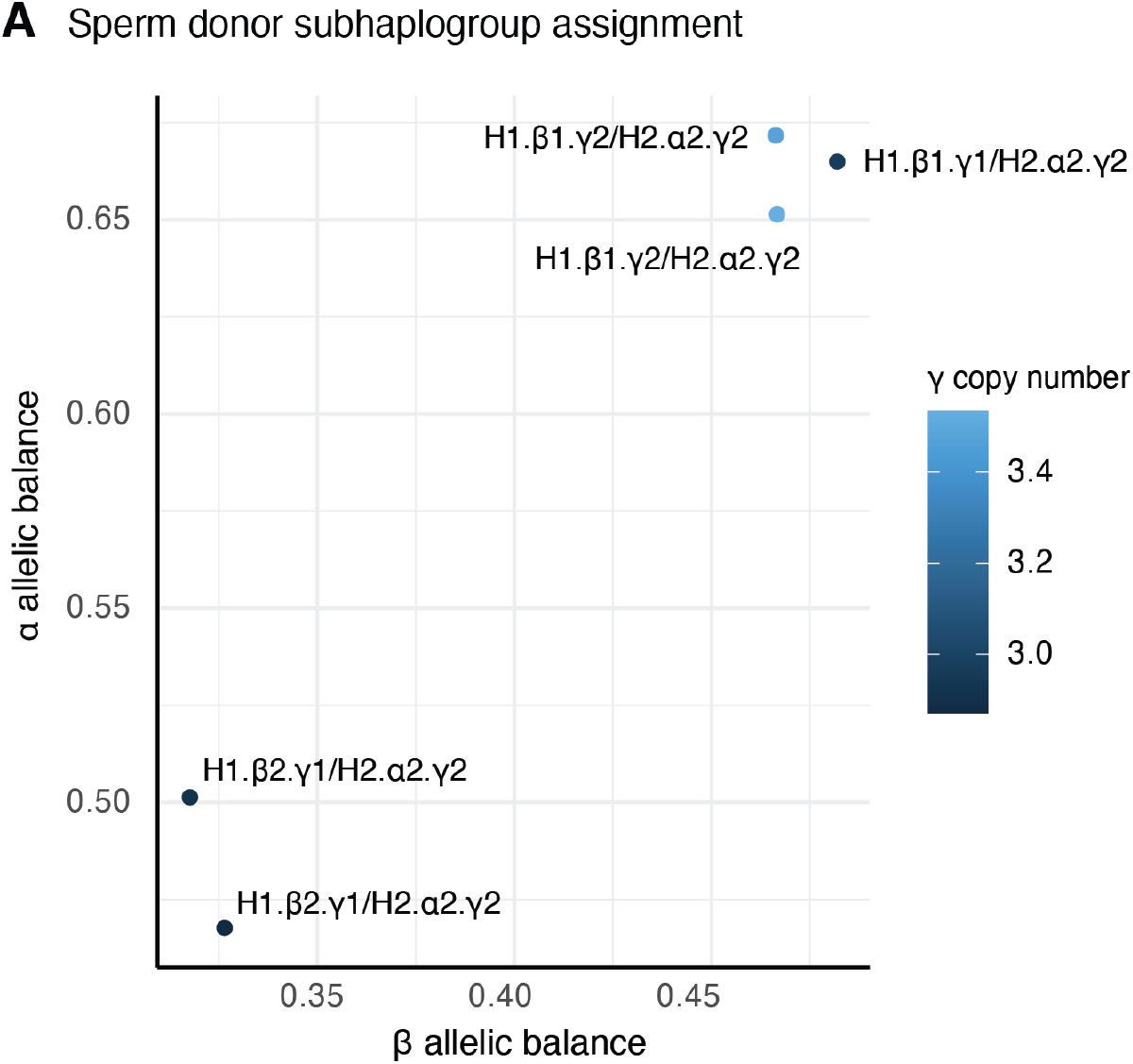
Subhaplogroup assignment in H1/H2 sperm donors. **A)** Allelic balance at α and β’ regions for the 5 H1/H2 sperm donors, colored by diploid copy number of the γ duplication. Each donor is labeled by assigned subhaplogroup based on allelic balance of α and β’ and copy number of α, β, γ.

**Supplemental Figure 3.**
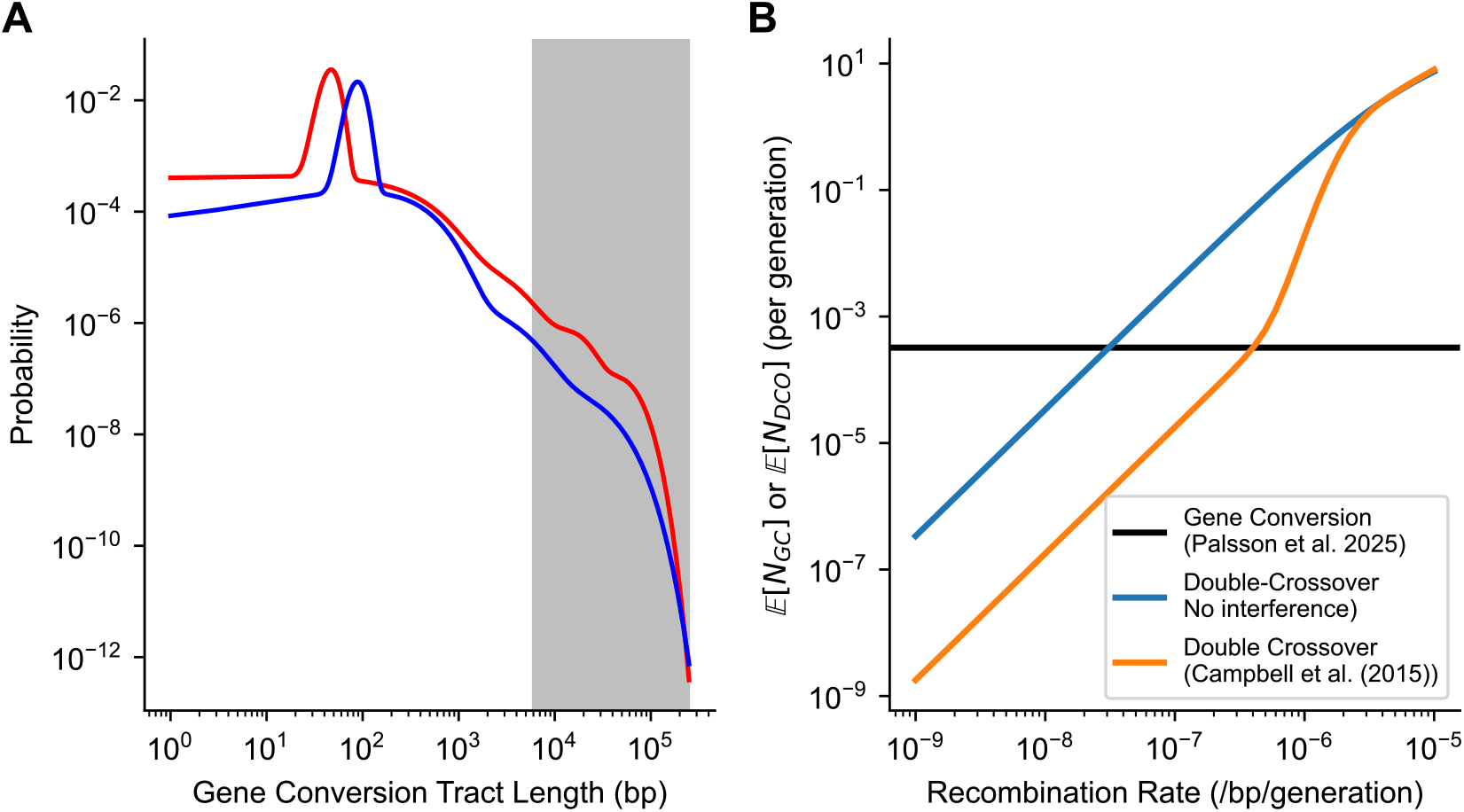
Comparison of gene conversion versus double crossovers for observed two-switch tract lengths. **A)** Length distribution of gene conversions observed in Palsson *et al*. 2025^39^. Tract lengths are modeled as a mixture of negative binomial distributions (see Supplemental Text), and shaded region reflects length scales of H1/H2 tracts observed in the inversion region. **B)** Expected number of gene conversion tracts or double-crossover events within a 478 kb region (restricting to ~5-250 kb tracts). With higher assumed recombination rates within the region, double crossovers can become as common as gene conversion tracts of this size. However, for recombination rates *O*(10^−8^) and realistic interference, we observe that the ratio of gene conversions to double crossovers is ~1628:1, suggesting that these tracts are the result of gene conversion events.

**Supplemental Figure 4.**
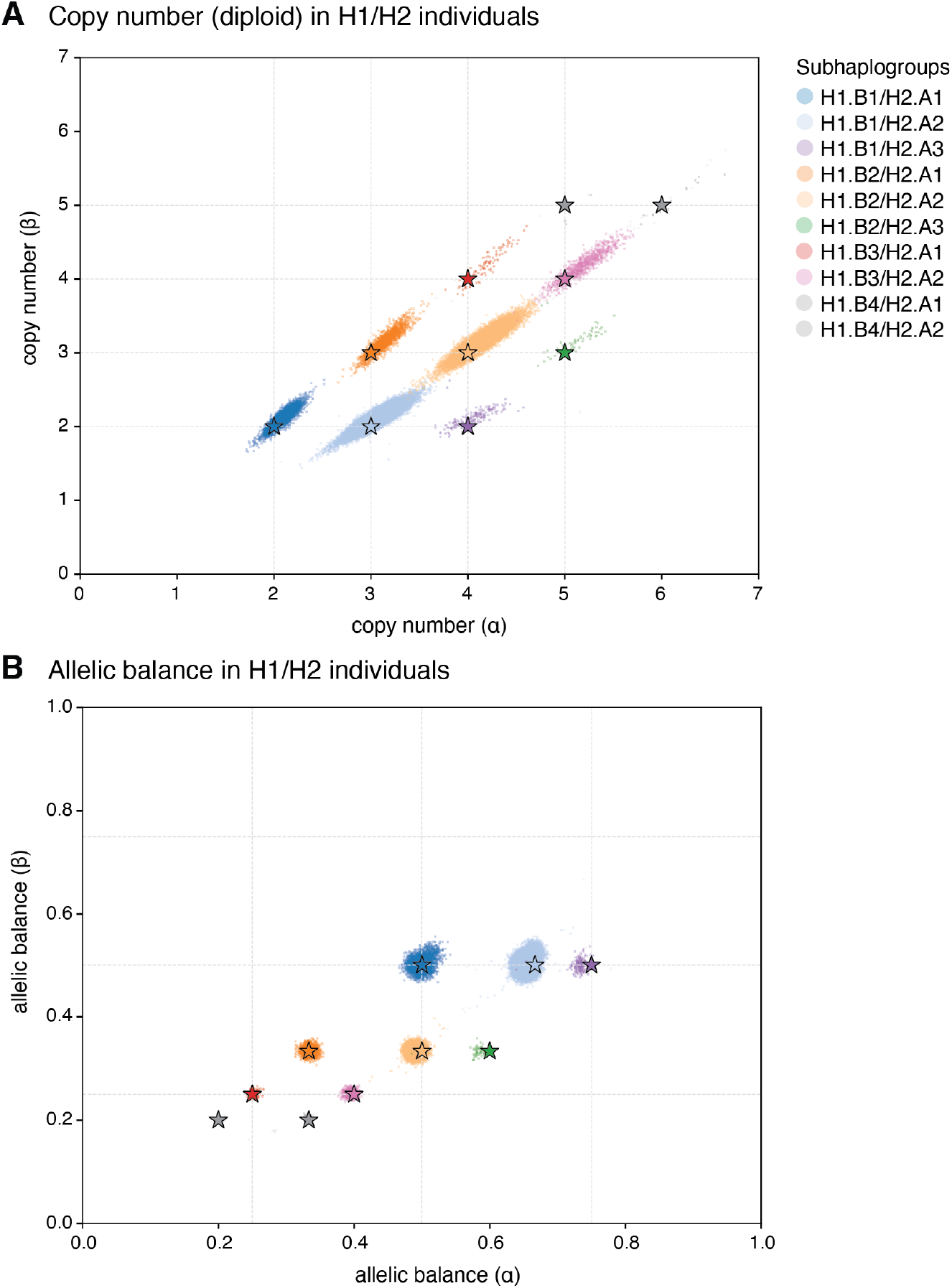
Subhaplogroup assignment in H1/H2 All of Us participants. **A)** Copy number (diploid) at α and β’ regions for the H1/H2 All of Us samples. Stars are placed at centroids for possible copy number combinations; samples are colored by assigned subhaplogroups based on closest centroids (clustering performed on copy number and allelic balance). **B)** Allelic balance at α and β’ regions (mean allelic balance at H1-H2 diagnostic SNPs). Stars are placed at centroids for possible allelic balance combinations; samples are colored by assigned subhaplogroups.

**Supplemental Figure 5.**
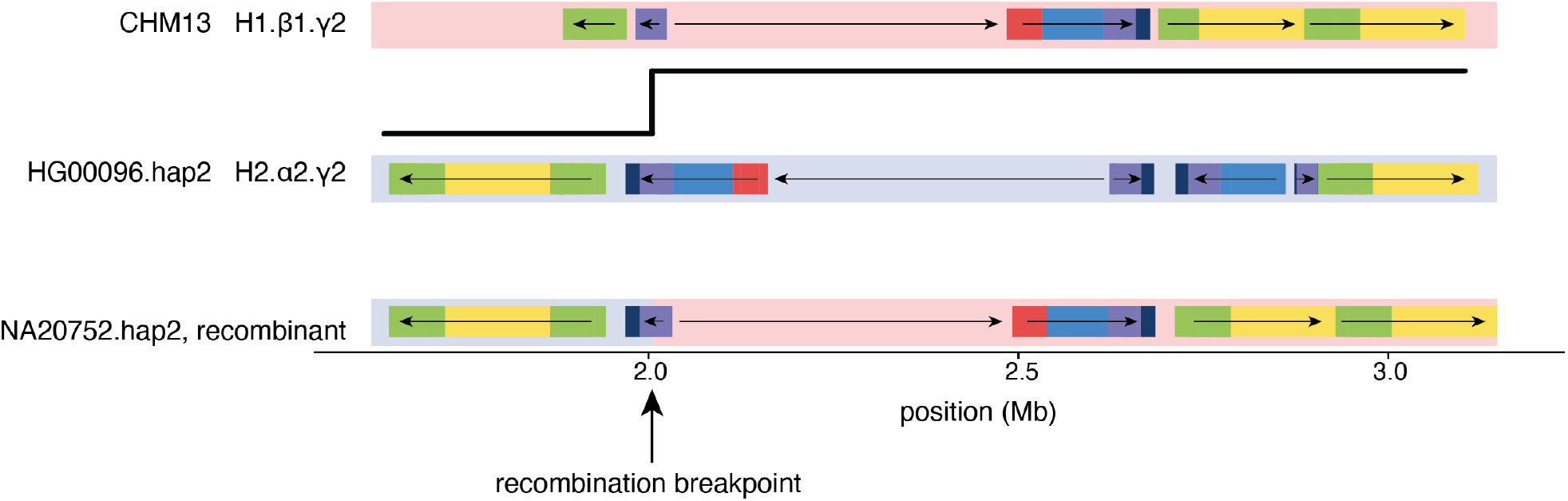
Haplotype-resolved genome assembly with H1-H2 recombination within SDs. Location of SDs in a representative assembly for H1.β1.γ2 (CHM13), H2.α2.γ2 (HG00096.hap2) and a recombinant haplotype (NA20752.hap2). The location of the recombination breakpoint in assembly NA20752.hap2 is highlighted, with a schematic (black line) of the likely non-allelic homologous recombination event between H2 and H1 haplotypes that created this haplotype. H1 sequence is highlighted in light pink, H2 in light blue.

**Supplemental Figure 6.**
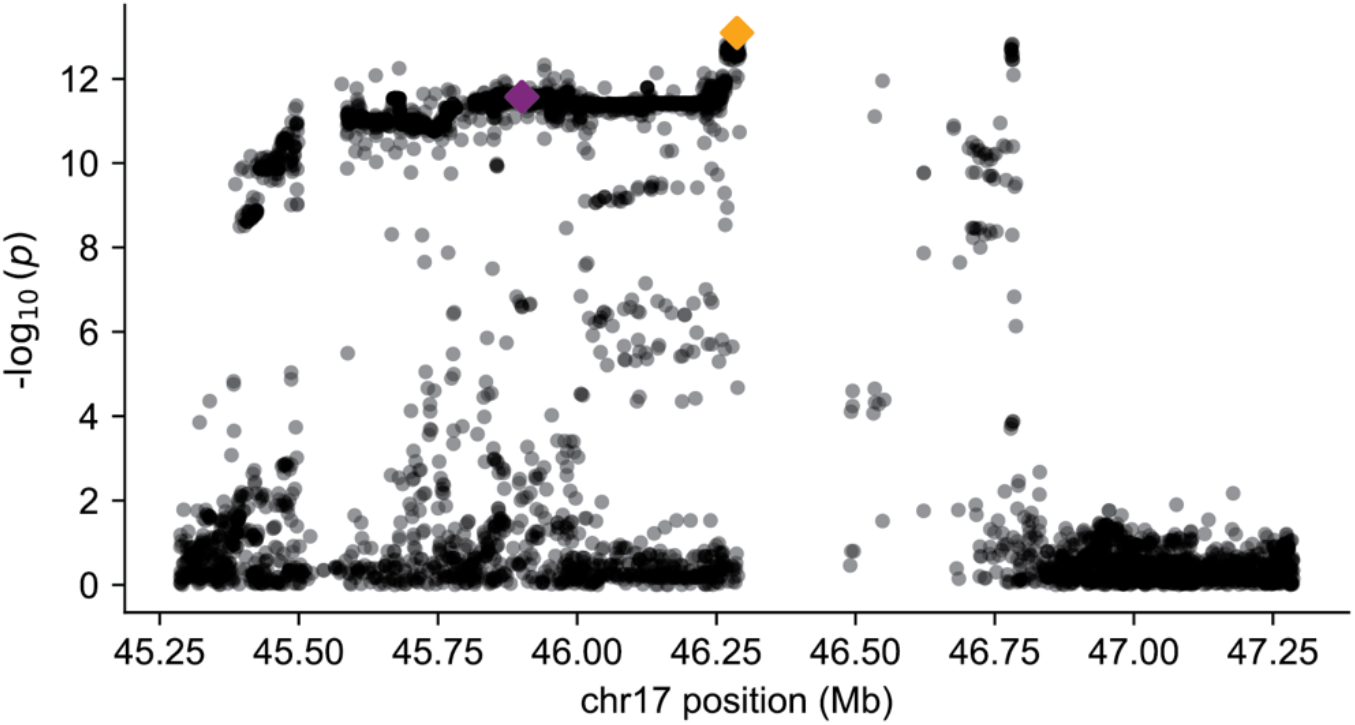
Locus association with maternal recombination at 17q21.31 in pre-implantation embryos. Association of variants with maternal recombination rate from ^43^. The orange diamond indicates the lead variant, chr17:46287153:C:G, and the purple diamond indicates H1-H2 tag SNP, chr17:45900675:C:T, used in this study.

**Supplemental Figure 7.**
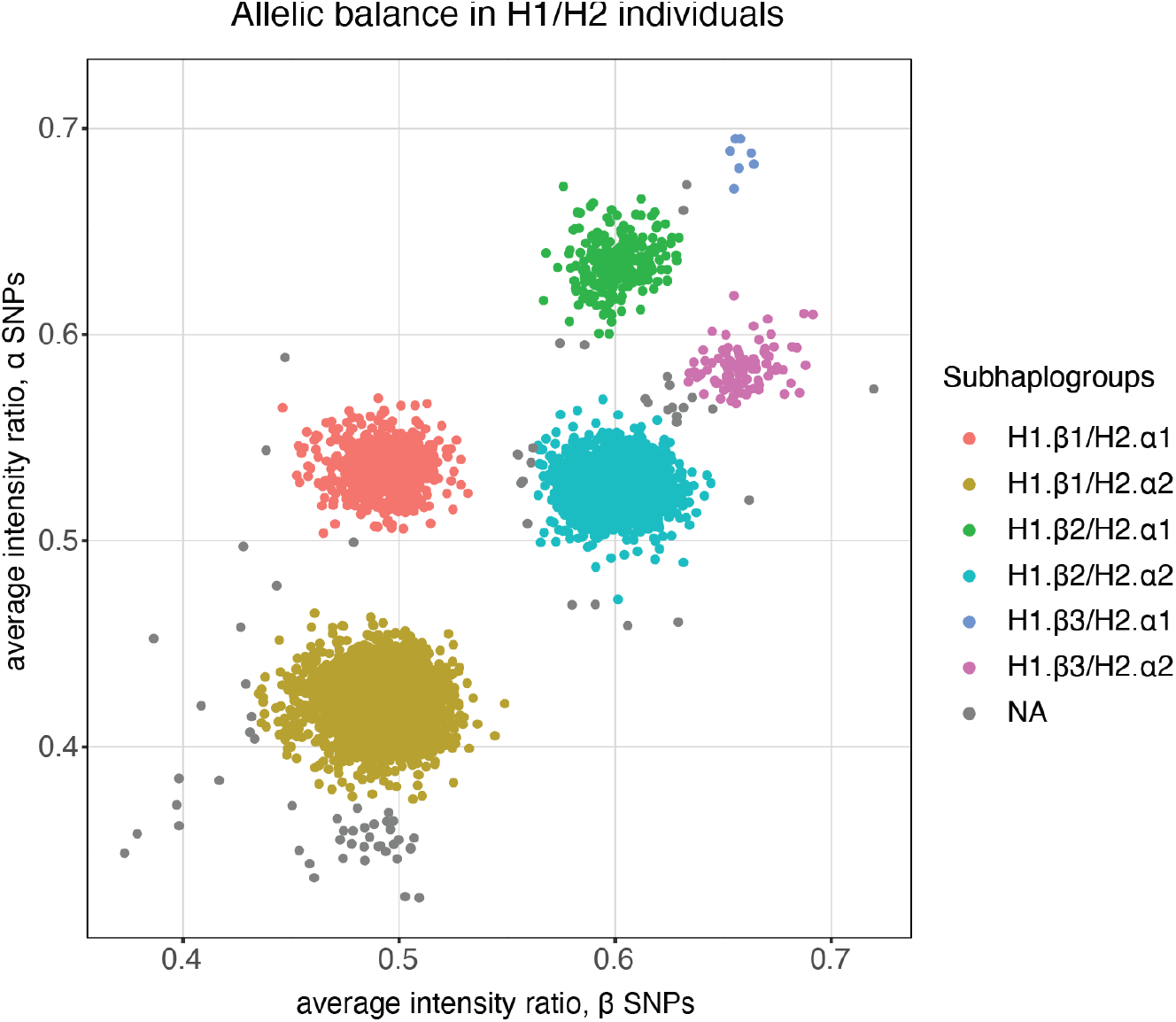
Subhaplogroup assignment in H1/H2 recombination data samples. Average intensity ratio, B/(A+B) where A and B are raw array probe intensities from H2 and H1 alleles respectively, for α SNPs (rs2732589, rs1918792, rs2732631, rs2668692) and β’ SNPs (rs17660907, rs4383188) from parent individuals in a parent-embryo recombination dataset ^43^. Subhaplogroups are assigned based on allelic balance of α and β’ SNPs, inferred from average intensity ratios.

**Supplemental Figure 8.**
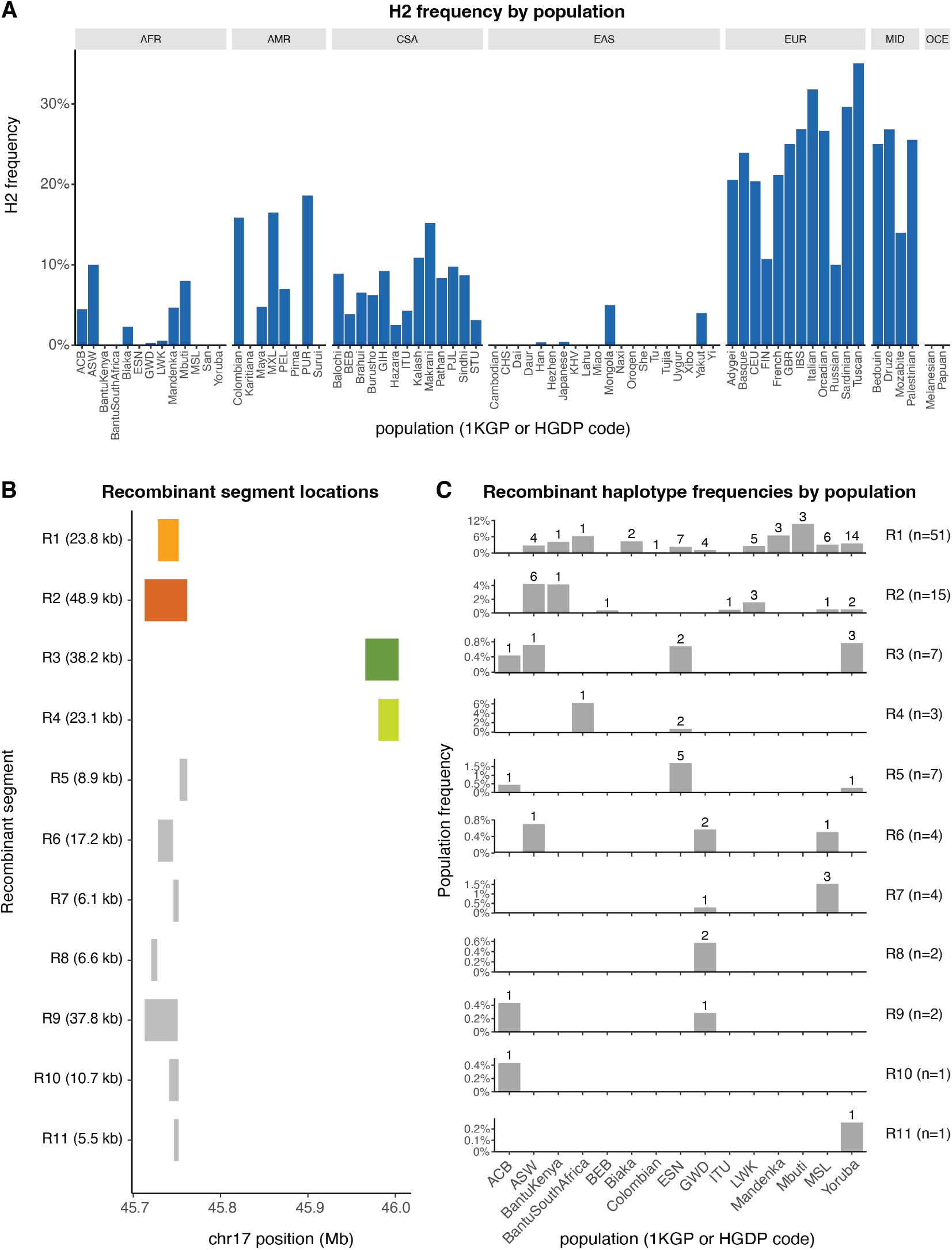
Inversion and recombinant haplotype frequencies in the 1KGP and HGDP dataset. **A)** Population frequencies of H2 by 1KGP and HGDP population code. **B)** Location of recombinant segments found within chr17:45.6-46.1 Mb. The four segments analyzed in detail are colored. Segment lengths are provided in parentheses. **C)** The population frequencies of each recombinant segment by 1KGP and HGDP population code. Number of recombinant haplotypes per population are listed above bar plot, with total number of recombinant haplotypes across the dataset provided in parentheses.

**Supplemental Figure 9.**
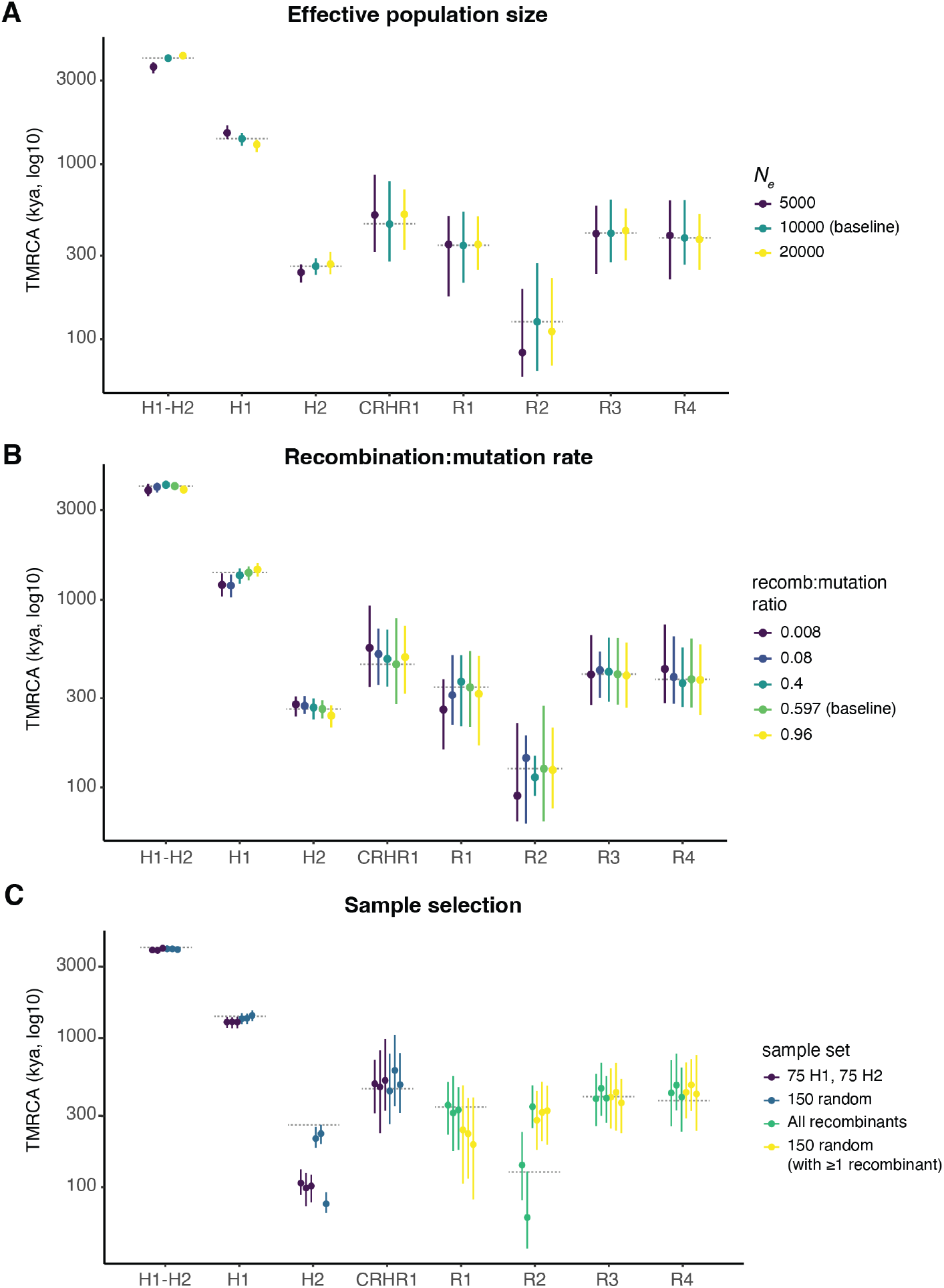
Sensitivity of coalescent time estimates with ARGs to input parameters and sample selection. TMRCA estimates for events of interest, varying effective population size (*N*_*e*_) as 5,000, 10,000 (baseline value) and 20,000 (**A**), varying the ratio of recombination to mutation rate as 0.008, 0.08, 0.4, 0.597 (baseline value) and 0.96 (**B**), and selecting different haplotype panels for making the ARGs (**C**). Panels of different sample sets tested include a random selection of 75 H1 and 75 H2 haplotypes, a random selection of 150 haplotypes, a selection of all recombinants (for segment of interest) plus random haplotypes to reach 150 total, and a random selection of 150 haplotypes (requiring that at least 1 recombinant haplotype for the segment of interest is included), with all haplotypes selected from the HGDP and 1KGP vcf. Three independent runs are shown per panel type. Points indicate posterior median, with 95% credible interval from ARGs posterior shown as lines. Dashed gray lines indicate estimates reported in Figure 5.

**Supplemental Figure 10.**
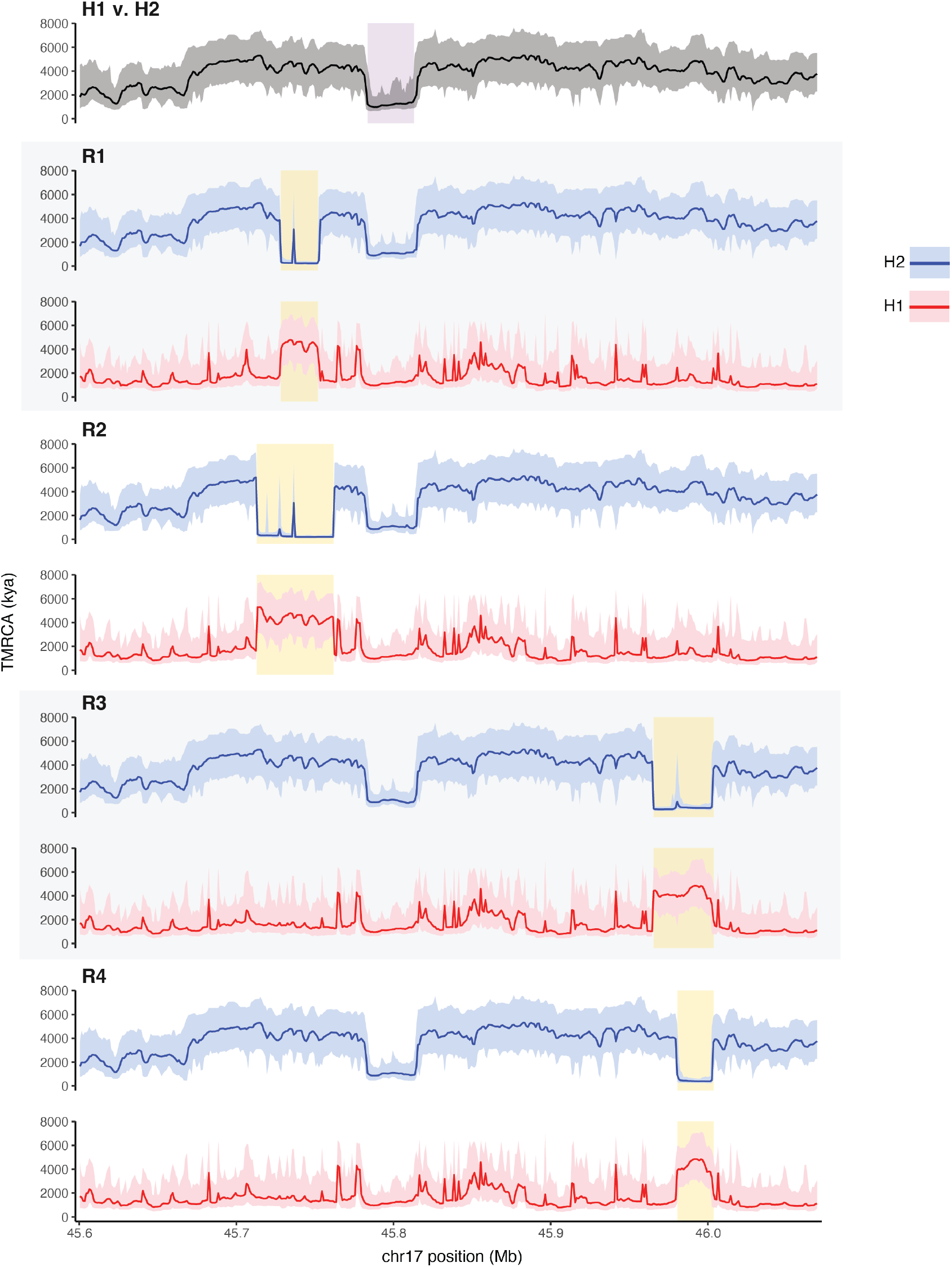
Windowed coalescence times across the inversion region. Divergence time estimates in 1-kb windows for H1 versus H2 haplotypes, and each set of recombinant haplotypes versus H2 (blue) and H1 (red). The median TMRCA estimate is shown as a solid line, with 95% credible intervals per window shaded. The *CRHR1* recombination region is highlighted in purple, and each recombinant segment is highlighted in yellow on its respective plot.

**Supplemental Figure 11.**
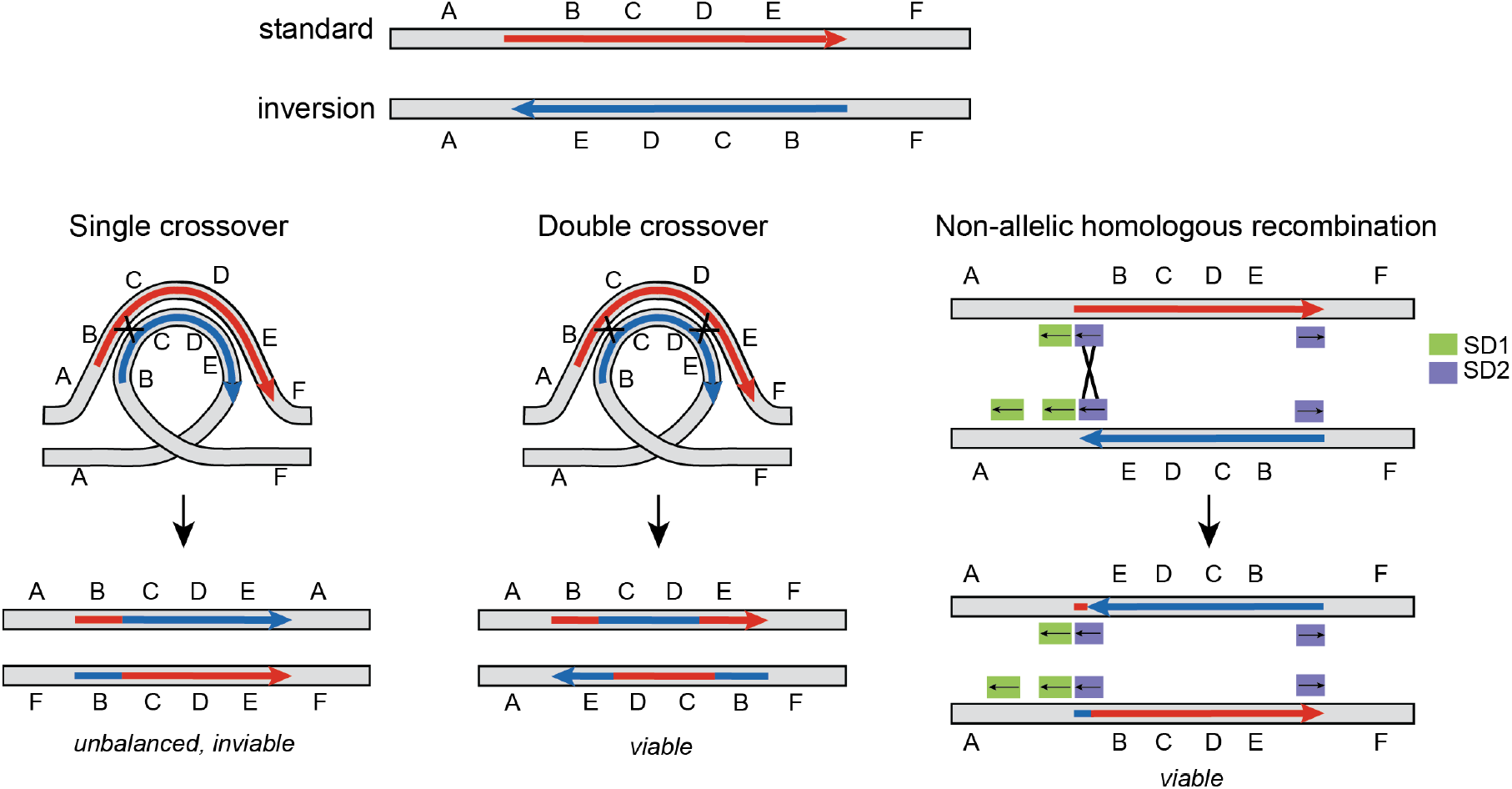
Mechanisms of gene flux at inversions. Standard and inversion arrangements can align during meiosis via an inversion loop. A single-crossover within the inversion loop yields unbalanced gametes (major chromosomal loss and gain). Double crossovers within the inversion loop yield balanced, viable gametes. NAHR between co-linear segmental duplications at an inversion breakpoint yields viable gametes without major chromosomal loss or gain.

**Supplemental Table 1.** Summary of inversion recombination rates in sperm donors. Per-donor counts of callable sperm cells, single crossovers and two-switch events within the inversion for H1/H2 donors.

| Subhaplogroup | # of callable sperm | # of single crossovers | # of two-switch events |
| --- | --- | --- | --- |
| H1.β2.g1/H2.α2.g2 | 1,644 | 0 | 3 |
| H1.β2.g1/H2.α2.g2 | 1,706 | 2 | 2 |
| H1.β1.g2/H2.α2.g2 | 1,267 | 1 | 4 |
| H1.β1.g2/H2.α2.g2 | 1,652 | 7 | 1 |
| H1.β1.g1/H2.α2.g2 | 1,898 | 7 | 3 |

**Supplemental Table 2.** Summary of All of Us recombinant haplotypes. Haplotype location (segment start, segment end, in Mb); number, ancestries and majority ancestry of samples carrying the haplotype; and subhaplogroup on which the recombinant segment resides. If haploid copy number could not be determined, possible subhaplogroups are listed. β(H1) denotes β copy from H1, β(H2) denotes β copy from H2 haplotype, α_p_denotes partial α copy, β_p_denotes partial β copy.

| Seg. ID | Start (Mb) | End (Mb) | # Samples | Ancestries | Majority ancestry | Subhaplogroup |
| --- | --- | --- | --- | --- | --- | --- |
| | 45.668 | 45.763 | <=20 | eur | eur | β1 |
| | 45.701 | 45.766 | <=20 | eur,sas | eur | β1,β1\|β2 |
| | 45.701 | 45.855 | <=20 | eur | eur | β1,β1\|β2 |
| | 45.705 | 45.718 | <=20 | eur | eur | β2 |
| | 45.707 | 45.771 | <=20 | amr,eur | amr | β1,β1\|β2 |
| | 45.709 | 45.724 | <=20 | eur | eur | β1 |
| R2 | 45.713 | 45.762 | >500 | afr,amr,eur,mid,sas | afr | β1,β1\|β2\|β3 |
| | 45.713 | 45.751 | 101-500 | afr,amr,eur | afr | β1,β1\|β2 |
| | 45.713 | 45.727 | 21-100 | afr,amr,eas,eur | afr | β1,β1\|β2 |
| | 45.713 | 45.96 | <=20 | eur | eur | β1 |
| | 45.713 | 45.763 | <=20 | afr | afr | β1 |
| | 45.721 | 45.762 | 21-100 | afr,amr,eur,sas | afr | β1,β1\|β2 |
| | 45.721 | 45.756 | <=20 | afr | afr | β1 |
| R1 | 45.728 | 45.752 | >500 | afr,amr,eur,mid,sas | afr | β1,β1\|β2\|β3 |
| | 45.728 | 45.745 | 21-100 | afr,amr | afr | β1,β1\|β2 |
| | 45.728 | 45.753 | <=20 | afr | afr | β1 |
| | 45.728 | 45.762 | <=20 | afr | afr | β1 |
| | 45.728 | 45.737 | <=20 | mid | mid | β1 |
| | 45.733 | 45.752 | <=20 | afr | afr | β1 |
| | 45.741 | 45.752 | 21-100 | afr,amr | afr | β1,β1\|β2 |
| | 45.746 | 45.752 | 21-100 | afr,amr,eur | amr | β1,β2 |
| | 45.752 | 45.762 | <=20 | afr,eur | afr | β1 |
| | 45.753 | 45.762 | 21-100 | afr,amr,eur | afr | β1 |
| | 45.763 | 45.96 | <=20 | amr,eur | eur | β1,β1\|β2 |
| | 45.763 | 45.941 | <=20 | eur | eur | β2 |
| | 45.766 | 45.855 | <=20 | eur | eur | β1 |
| | 45.767 | 46.006 | <=20 | eur | eur | β1\|β2 |
| | 45.775 | 45.855 | <=20 | afr | afr | β1 |
| | 45.795 | 45.856 | 21-100 | afr,amr,eur | eur | β1,β1\|β2\|β3 |
| | 45.795 | 45.96 | <=20 | eur,sas | eur | β1,β2 |
| | 45.795 | 45.844 | <=20 | eur | eur | β2 |
| | 45.797 | 45.858 | <=20 | sas | sas | β1 |
| | 45.81 | 45.96 | 101-500 | afr,amr,eur | eur | β1,β1\|β2\|β3 |
| | 45.81 | 45.855 | <=20 | eur | eur | β1,β1\|β2 |
| | 45.81 | 45.941 | <=20 | eur | eur | β1 |
| | 45.81 | 45.829 | <=20 | eur | eur | β1\|β2 |
| | 45.81 | 45.872 | <=20 | eur | eur | β1 |
| | 45.81 | 45.916 | <=20 | eur | eur | β1\|β2 |
| | 45.848 | 45.91 | <=20 | eur | eur | β1 |
| | 45.856 | 45.941 | <=20 | amr,eur | eur | β1,β1\|β2\|β3 |
| | 45.856 | 45.869 | <=20 | eur | eur | β1 |
| | 45.856 | 45.944 | <=20 | eur | eur | β1 |
| | 45.856 | 45.867 | <=20 | eur | eur | β1 |
| | 45.856 | 46.003 | <=20 | eur | eur | β1\|β2 |
| | 45.856 | 46.019 | <=20 | eur | eur | β1\|β2 |
| | 45.856 | 45.986 | <=20 | eur | eur | β1\|β2\|β3 |
| | 45.856 | 45.96 | <=20 | eur | eur | β1,β1\|β2\|β3 |
| | 45.868 | 45.96 | <=20 | eur | eur | β1 |
| | 45.869 | 45.941 | <=20 | eur | eur | β1 |
| | 45.917 | 45.941 | <=20 | eur | eur | β1 |
| | 45.942 | 45.96 | <=20 | eur | eur | β1\|β2\|β3 |
| R3 | 45.966 | 46.004 | >500 | afr,amr,eur,sas | afr | β1,β1\|β2\|β3 |
| | 45.966 | 46.002 | <=20 | afr | afr | β1,β1\|β2 |
| R4 | 45.981 | 46.004 | 101-500 | afr,amr,eur | afr | β1,β1\|β2 |
| | 45.981 | 45.996 | 21-100 | afr | afr | β1 |
| | 45.983 | 46.004 | <=20 | afr,amr | afr | β1 |
| | 46.006 | 46.029 | <=20 | eur | eur | β1 |
| | 46.012 | 46.027 | <=20 | eur | eur | β1 |
| | 45.659 | 45.795 | <=20 | eur | eur | α2 |
| | 45.717 | 45.726 | <=20 | eur | eur | α2 |
| | 45.762 | 45.784 | <=20 | eur | eur | α2 |
| | 45.766 | 45.98 | <=20 | amr | amr | α2 |
| | 45.766 | 45.808 | <=20 | afr | afr | α1 |
| | 45.766 | 45.778 | <=20 | eur | eur | α2 |
| | 45.767 | 45.774 | <=20 | amr | amr | α2 |
| | 45.795 | 45.874 | <=20 | amr | amr | α2 |
| | 45.795 | 45.819 | <=20 | afr | afr | α2 |
| | 45.81 | 45.83 | <=20 | eur | eur | $\alpha_2$ |
| | 45.815 | 45.841 | <=20 | eur | eur | $\alpha_2$ |
| | 45.832 | 45.867 | <=20 | eur | eur | $\alpha_2$ |
| | 45.961 | 45.976 | <=20 | amr | amr | $\alpha_2$ |
| | 46.011 | 46.022 | <=20 | eur | eur | $\alpha_2$ |
| | 46.071 | 46.092 | <=20 | mid | mid | $\alpha_2$ |
| | 45.856 | unknown | <=20 | eur | eur | $\beta(H_2)\beta(H_1)$ |
| | 45.961 | unknown | <=20 | eur | eur | $\beta(H_2)\beta(H_1)$ |
| | 46.086 | 46.163 | <=20 | eur | eur | $\beta_p(H_2)\alpha_p(H_1)$ |
| | 45.666 | unknown | <=20 | eur | eur | $\alpha(H_2)\beta(H_1)$ |
| | 45.68 | unknown | <=20 | eur | eur | $\alpha(H_2)\beta(H_1)$ |
| | 45.701 | unknown | 21-100 | afr,amr,eur | eur | $\alpha(H_2)\beta(H_1)$ |
| | 45.705 | unknown | 21-100 | afr,amr,eur | eur | $\alpha(H_2)\beta(H_1)$ |
| | 45.762 | unknown | <=20 | afr,eur | afr | $\alpha(H_2)\beta(H_1)$ |
| | 45.763 | unknown | 21-100 | amr,eur | eur | $\alpha(H_2)\beta(H_1)$ |
| | 45.771 | unknown | 21-100 | afr,amr,eur,sas | eur | $\alpha(H_2)\beta(H_1)$ |
| | 45.773 | unknown | <=20 | eur | eur | $\alpha(H_2)\beta(H_1)$ |
| | 45.784 | unknown | <=20 | amr,eur | eur | $\alpha(H_2)\beta(H_1)$ |
| | 45.81 | unknown | <=20 | eur | eur | $\alpha(H_2)\beta(H_1)$ |
| | 45.833 | unknown | <=20 | eur | eur | $\alpha(H_2)\beta(H_1)$ |
| | 45.856 | unknown | 21-100 | afr,amr,eur,mid,sas | eur | $\alpha(H_2)\beta(H_1)$ |
| | 45.858 | unknown | <=20 | eur | eur | $\alpha(H_2)\beta(H_1)$ |
| | 45.869 | unknown | 21-100 | afr,amr,eur | amr | $\alpha(H_2)\beta(H_1)$ |
| | 45.917 | unknown | <=20 | eur | eur | $\alpha(H_2)\beta(H_1)$ |
| | 45.942 | unknown | <=20 | eur,mid | eur | $\alpha(H_2)\beta(H_1)$ |
| | 45.961 | unknown | 21-100 | afr,eur | eur | $\alpha(H_2)\beta(H_1)$ |
| | 45.986 | unknown | <=20 | amr | amr | $\alpha(H_2)\beta(H_1)$ |
| | 46.021 | unknown | <=20 | eur | eur | $\alpha(H_2)\beta(H_1)$ |
| | 46.107 | unknown | <=20 | afr,eur | eur | $\beta_p(H_2)\beta(H_1)$ |
| | 46.136 | unknown | >500 | afr,amr,eas,eur,mid,sas | eur | $\alpha(H_2)\beta(H_1)$ |
| | 46.169 | unknown | <=20 | eur | eur | $\alpha_p(H_2)\beta(H_1)$ |
| | 46.173 | unknown | <=20 | afr | afr | $\alpha_p(H_2)\beta(H_1)$ |

**Supplemental Table 3.** Haplotype-specific effects on maternal crossover rates. All effects are estimated using a Welch’s t-test (two-sided) relative to the H1/H1 haplotype assignments.

| <u>Category</u> | <u>mu null</u> | <u>mu alt</u> | <u>n alt</u> | <u>p value</u> | <u>df</u> |
| --- | --- | --- | --- | --- | --- |
| H2/H2 | -0.059839 | 0.281488 | 385 | 4.4925e-11 | 428 |
| H1.β1/H2.α1 | -0.059839 | 0.132295 | 180 | 0.012662 | 187 |
| H1.β1/H2.α2 | -0.059839 | 0.111525 | 1863 | 4.1646e-11 | 2888 |
| H1.β2/H2.α1 | -0.059839 | 0.23436 | 58 | 0.014931 | 58 |
| H1.β2/H2.α2 | -0.059839 | 0.105905 | 682 | 0.000039 | 812 |
| H1.β1/H1.α(H2)β(H1) | -0.059839 | -0.072945 | 17 | 0.960894 | 16 |

## Supplemental Text

We observed multiple tracts of H2 ancestry on H1 background (and vice-versa) at the chr17q21.31 inversion. Here we try to reason whether these tracts may be the result of gene conversion events or double crossovers based on estimates of complete recombination maps in human pedigrees and parameters of crossover interference^39,41^.

### Models

#### Shared Notation

The central goal of our analysis is to evaluate the probability of tracts of a specified length scale on the H1 or H2 inversion backgrounds as being a result of gene conversion or a double crossover event. There are several parameters that are shared between the models for gene conversion and double crossover respectively which are detailed below.

| Variable | Interpretation | Units |
| --- | --- | --- |
| $r$ | Recombination/crossover rate or gene conversion rate | per basepair, per generation |
| $L$ | Length of the chromosomal region affected | basepairs |
| $\nu$ | Crossover interference parameter in Houseworth-Stahl Model | NA |
| $a, b$ | Boundaries of tract length covered | basepairs |
| $p$ | Fraction of crossovers which “escape” interference | NA |

#### Gene Conversion

Let us suppose that gene conversion occurs at a rate *r* per-basepair per generation and the tract length of gene conversion is based on an arbitrary distribution with cumulative mass function *F*^*∗*^(*x*)). In a region of size *L*, we therefore have the following probability for a gene conversion tract in a generation to be within a specific size range *l* ∈ *(a, b)*:

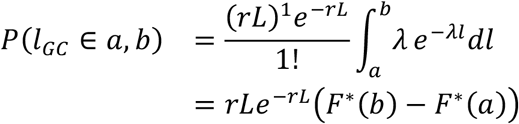

It may also be easier to consider this as the expected number of gene conversion events in a generation within the range of (*a,b*) as:

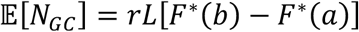

This notation is particularly helpful when segments are modeled as a mixture of distributions as shown in recent work (Palsson et al., 2025).

#### Double Crossover

For a tract to be the result of a double-crossover and within a certain length scale, we need to first condition on the probability of exactly two crossovers forming in a region of length *G* Morgans, where *G* = *rL*. The occurrence of crossovers can be approximated as a Poisson distributed random variable within the region. One final component is the expected “gap” *l* (in Morgans) between two crossovers — which is modeled as a mixture of events with and without crossover interference.

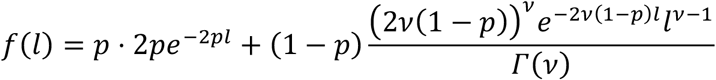

However, unlike the model for gene conversion (which only needs to have one occurrence in the interval from [0, *L*]) we need to require that two crossover events occur within [0, *G*]. We can broadly conceptualize this as some number of crossovers placed on a very long region and artificially placing a smaller window *G* within this interval and ask if it captures a specific gap *l*. Assuming that the intensity of crossovers is one crossover per-Morgan (*G* is in morgans), we can define:

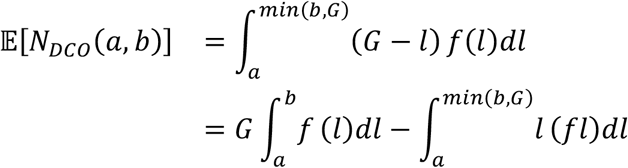

The second term is a term which accounts for the fact that the boundary at *G* can be affected by longer gaps (which will not be counted). Integrating this, we obtain the following expression:

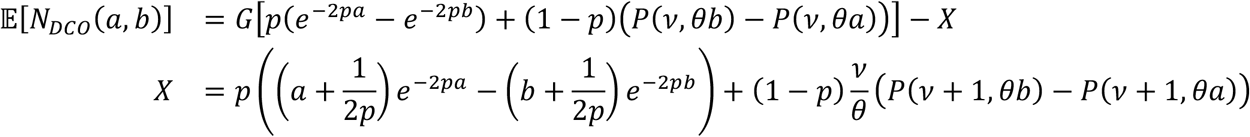

In the above equation, *P*(*⋅*) is the normalized incomplete gamma function, θ = *2v*(1 − *p*) and *r* is the recombination rate. If *G* ≫ *b*, then the second portion of the equation may be dropped, but in the case of this inversion, that is not the case so we will proceed with the exact solution here. To translate these to the physical length-scale, we will scale the variables *a, b* by a recombination rate *r* (and *G* = *rL* already).

### Empirical Parameters

We model gene conversion tracts the same as in recent work as a multi-component mixture of Negative binomial tract-length distributions, with initiation treated as a Poisson process along the genome^39^. The parameters below are directly extracted from Table S4 in Palsson *et al*. 2025 and shown here in a summarized form^39^. The probability mass for each length-scale is shown across both sexes, with maternal gene conversions having higher mass within the larger segments relevant to observations in the chr17q21.31 region (Supplemental Figure 3A).

#### Paternal NCOs

| Component | Class | Percent NCOs | Mean NCO length | Sigma |
| --- | --- | --- | --- | --- |
| 1 | Short | 87.77 | 91 bp | 16.6 |
| 2 | Short | 11.09 | 374 bp | 325 |
| 3 | Extended | 0.89 | 2.9 kb | 2.9 |
| 4 | Extended | 0.24 | 23.2 kb | 21.6 |

#### Crossover interference (Gamma-escape / Housworth–Stahl model)

Inter-crossover spacing modeled as a renewal process with a two-component mixture: an “escape”/no-interference component (Exponential) and an “interfering” component (Gamma, shape *v*). These sex-specific estimates are based on whole-genome parameter estimates^41^.

| Parameter | Maternal | Paternal | Description |
| --- | --- | --- | --- |
| nu ($\nu$) | 7.19 | 8.93 | Interference strength (gamma shape) |
| p | 0.078 | 0.067 | Escape-from-interference probability |

### Results

We can compare the expected number of gene conversions or double crossovers within the approximately 478 kb single-copy region of the inversion over a generation. We restricted to segments which are between ~5-250 kb to reflect the tract lengths observed in sperm-sequencing and AllofUs individuals at this locus. Comparing these relatively under observed parameters for the gene conversion and crossover process respectively identifies the relative abundance between the two classes and suggests that for genome-wide average recombination rates in the region that gene conversions are substantially more likely. For example, the expected number of gene conversion tracts is ~2.9 × 10^−4^ per generation, whereas with realistic interference parameters and a recombination rate *r* = 10^−8^ there are 1.79 × 10^−7^ expected double crossovers per generation (Supplemental Figure 3B).

